# Floral scents of a deceptive plant are hyperdiverse and under population-specific phenotypic selection

**DOI:** 10.1101/2021.04.28.441155

**Authors:** Gfrerer Eva, Laina Danae, Gibernau Marc, Fuchs Roman, Happ Martin, Tolasch Till, Trutschnig Wolfgang, Anja C. Hörger, Comes Hans Peter, Dötterl Stefan

## Abstract

Floral scent is a key mediator in plant-pollinator interactions; however, little is known to what extent intraspecific scent variation is shaped by phenotypic selection, with no information yet in deceptive plants. We recorded 291 scent compounds in deceptive moth fly-*pollinated Arum maculatum* from various populations north vs. south of the Alps, the highest number so far reported in a single plant species. Scent and fruit set differed between regions, and some, but not all differences in scent could be explained by differential phenotypic selection in northern vs. southern populations. Our study is the first to provide evidence that phenotypic selection is involved in shaping geographic patterns of floral scent in deceptive plants. The hyperdiverse scent of *A. maculatum* might result from the plant’s imitation of various brood substrates of its pollinators.

## INTRODUCTION

About 88% of angiosperms are cross-pollinated by animals^1^ that are attracted to flowers by multifaceted cues^2^. Together with visual cues, the main attractant for pollinators is floral scent^3,4^. Therefore, scent has strong effects on pollinator visitation and frequency, and hence the plant’s reproductive success^3,5^. With more than 2,000 floral volatile organic compounds (VOCs) described^4,6^, and an average of 20–60 VOCs per species^7^, floral scent blends can tremendously vary among species in terms of composition and quantity. Consequently, they facilitate discrimination by pollinators among host plant species and contribute to reproductive isolation of closely related species^8–11^.

In addition to interspecific variation, floral scent is also known to vary intraspecifically, both within and among populations^5,12–16^. Such intraspecific variability might result directly from abiotic (*e*.*g*., temperature^17^, soil chemistry^18,19^) and/or biotic factors (*e*.*g*., herbivores^20,21^, microbes^22^). Given that scent is heritable^23–25^, intraspecific differences can also result from varying evolutionary forces, such as natural selection and genetic drift^4,26,27^. Although not explicitly demonstrated, genetic drift was suggested to be responsible for strong inter-population differences in floral scents^5^, or to counteract pollinator-mediated selection (in two *Yucca* species^28^). In contrast, natural selection on floral scent emission, both on total scent amount and on individual scent components, has been shown by analyses of phenotypic selection, correlating scent phenotypes and fitness measures^29–36^.

Phenotypic selection on floral scent can also vary intraspecifically, potentially leading to variable adaptive responses to spatially variable pollinator assemblages^32,36,37^. Until now, studies examining phenotypic selection on floral scent have tested rewarding, but not deceptive species, although the latter also often rely on luring and deceiving their pollinators with scents^38–40^. Compared to their rewarding relatives, non-rewarding species often display higher variation in scent and other traits attractive to pollinators^41–43^,and are frequently more pollen-limited^44–46^. In consequence, they might experience stronger selection on floral scent than rewarding species, as shown for floral traits other than scent^47^.

An ideal target for studying phenotypic selection on scent is the moth fly-pollinated and brood-site deceptive *Arum maculatum* L. (Araceae), which attracts its pollinators by olfactory deception. This perennial herb is widespread in Europe and shows high variation in fruit and seed sets within and among populations^48,49^. The main pollinators are two moth flies (*Psychoda phalaenoides* L. and *P. grisescens* Tonn., Psychodidae) that are attracted by the strong, dung-like inflorescence scent of *A. maculatum* while looking for oviposition sites and/or mating partners^50–52^. Previous analyses have shown that the scent profile of *A. maculatum* consists of up to 60 compounds, also differing among populations^53–58^. At least in part, this scent variation appears to reflect variation in the pollinator assemblages of *A. maculatum* across its distribution range^55,58^. In Central and much of Western Europe, high abundances of a single *Psychoda* species and sex (females of *P. phalaenoides*) were found^52^. In other regions (Mediterranean Europe and Western France), *A. maculatum* was visited in lower abundances but with a higher diversity of insects (mostly females of *P. phalaenoides* and both sexes of *P. grisescens*, plus a few other psychodid species and some other Diptera), also with some variation among populations^52^. This geographic pollinator variation is particularly pronounced north vs. south of the Alps^52, Laina, D. *et al*., unpubl.^ and matches a weak genetic (AFLP) subdivision of *A. maculatum* across this geographic barrier^59^. Presently, it is unclear whether those regional patterns of pollinator abundance/diversity and genetic variation between populations of *A. maculatum* are also reflected in their scent patterns. It is known, however, that the two main pollinating moth fly species have dissimilar floral scent preferences^55,58^. Hence, we assume that the dissimilar scent preferences of the two fly species, together with their different floral visitation in regions north vs. south of the Alps, could have led to differing selection pressures on scent among respective regional populations of *A. maculatum* from north vs. south of the Alps.

In this study, we investigated the floral scent characteristics and fruit set (as an indicator for female fitness) of *A. maculatum* in six populations north of the Alps vs. five populations south of the Alps and tested for phenotypic selection on scent in the largest and most extensively sampled population in each of the two regions. Specifically, we asked: (1) Do scent and fruit set differ between populations north vs. south of the Alps, and among populations within regions? (2) Is there phenotypic selection on floral scent in each of the most extensively sampled northern and southern population? And if so, (3) do compounds under selection differ between these two populations? Considering the differences in pollinator abundance and diversity between regions, and also among southern, but not northern populations^52^, we expect to find pronounced population differences in scent, both at the inter-regional level and within the southern region. When taking also the different olfactory preferences of pollinator species into account, we additionally expect lower fruit set south than north of the Alps, and different signs of selection in the most extensively sampled northern and southern populations.

## RESULTS

### Floral scent

The total absolute amount of scent was highly variable among the 233 sampled individuals (Fig. 1) of *A. maculatum* (range: 1−2,052 ng inflorescence^-1^ h^-1^; Table **1**). When taken together, northern plants released a three-fold lower amount of scent than those from the South, along with differences among populations within regions (permANOVA: *region*: pseudo-*F*_1,222_ = 25.70, *population* nested within *region*: pseudo-*F*_9,222_ = 5.36, both *P* < 0.001). For three of the five southern populations (MAH, MON, LIM), we estimated a median scent amount of *c*. 200 ng inflorescence^-1^ h^-1^, while DAO and UDI showed 1.5-fold higher and five-fold lower amounts, respectively. For three of the six northern populations (MUR, NEC, RÜM), corresponding median estimates ranged between 40 and 81 ng inflorescence^-1^ h^-1^, while in the remainder amounts were manifold higher (JOS and HOH) or lower (BUR) (Table **1**, Supporting Information Table **S1**).

**Fig 1.**
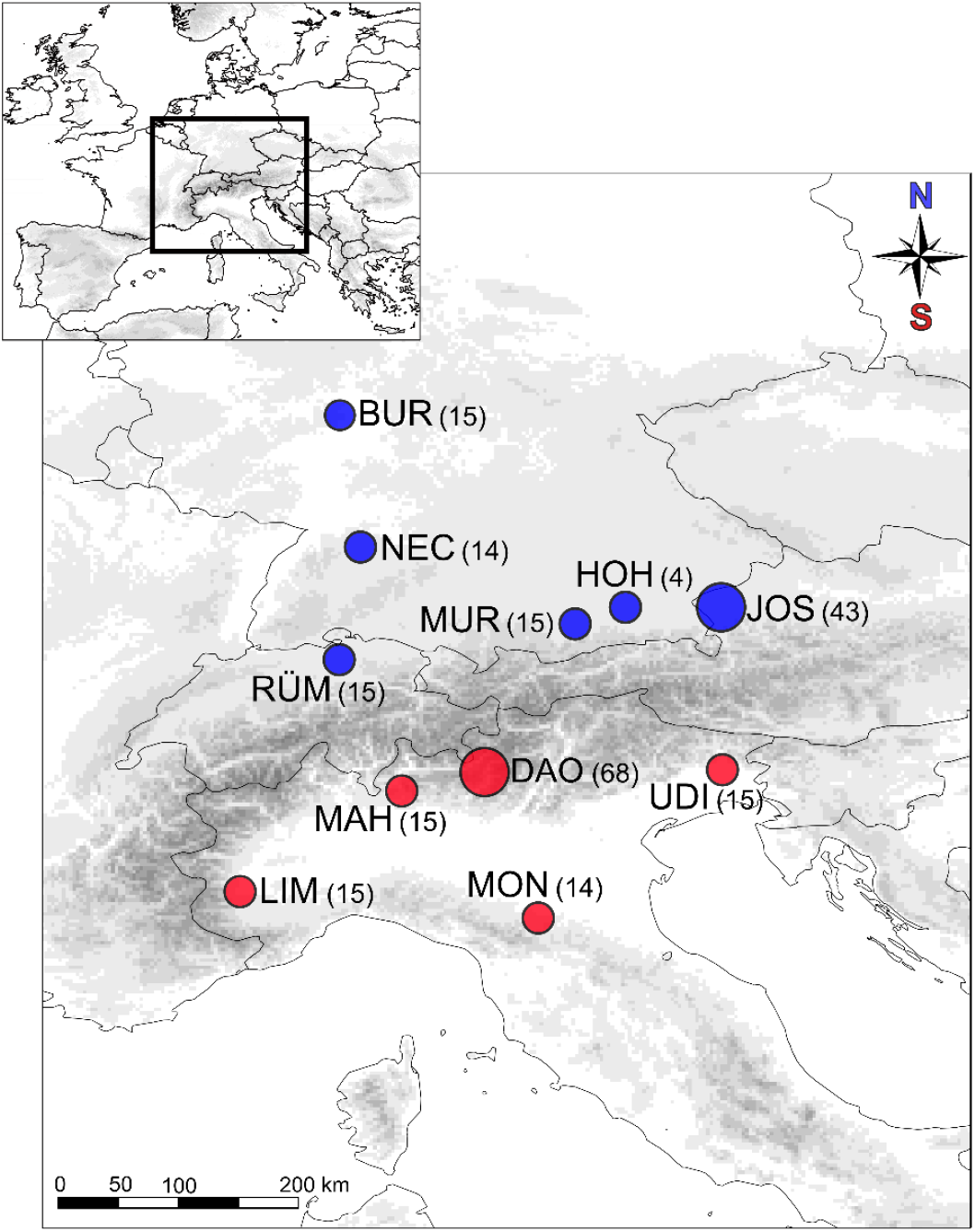
Sampling localities of *Arum maculatum* from north (blue) vs. south (red) of the Alps. Numbers in brackets give the number of individuals used for scent (and selection) analyses. The two most extensively sampled populations (JOS, DAO) are indicated by larger circles. *North*: JOS, Josefiau; BUR, Burg Hohenstein; HOH, Hohendilching; MUR, Murnau; NEC, Horb am Neckar; RÜM, Rümikon; *South*: DAO, Daone; LIM, Limone-Piemonte; MAH, Santa Maria Hoè; MON, Montese; UDI, Udine. The map was prepared using the ETOPO1 Global Relief Model^132^ and ArcGIS v.10.4 (ESRI, Redland, CA).

Across all scent samples, we detected a total of 291 floral volatiles (283 north vs. 265 south), and 92 of those could be chemically identified (Table **1**, and Supporting Information Table **S2**). A median of 102 compounds per individual was recorded (Fig. **2**), and the number of compounds was independent of the region (permANOVA: pseudo-*F*_1,222_ = 2.00, *P* = 0.15), but varied among populations within regions (pseudo-*F*_9,222_ = 4.57, *P* = 0.001). At the population level, between 166 (BUR) and 266 (JOS) compounds were recorded in the North, and between 88 (MON) and 254 (DAO) in the South (Fig. **2**). The two most extensively sampled northern (JOS) vs. southern (DAO) populations covered 92% vs. 87% of their respective regional diversity (Fig. **2**), and together 97% (283/291) of the total number of compounds (Table **1**, and Supporting Information Table **S2**). The five most frequent compounds, found in more than 99% of the samples, were the nitrogen-bearing compound indole, the monoterpenoids 3,7-dimethyloct-1-ene and *β*-citronellene, the sesquiterpenoid *β*-caryophyllene, and the unidentified UNK1492 (Table **1**).

**Fig 2.**
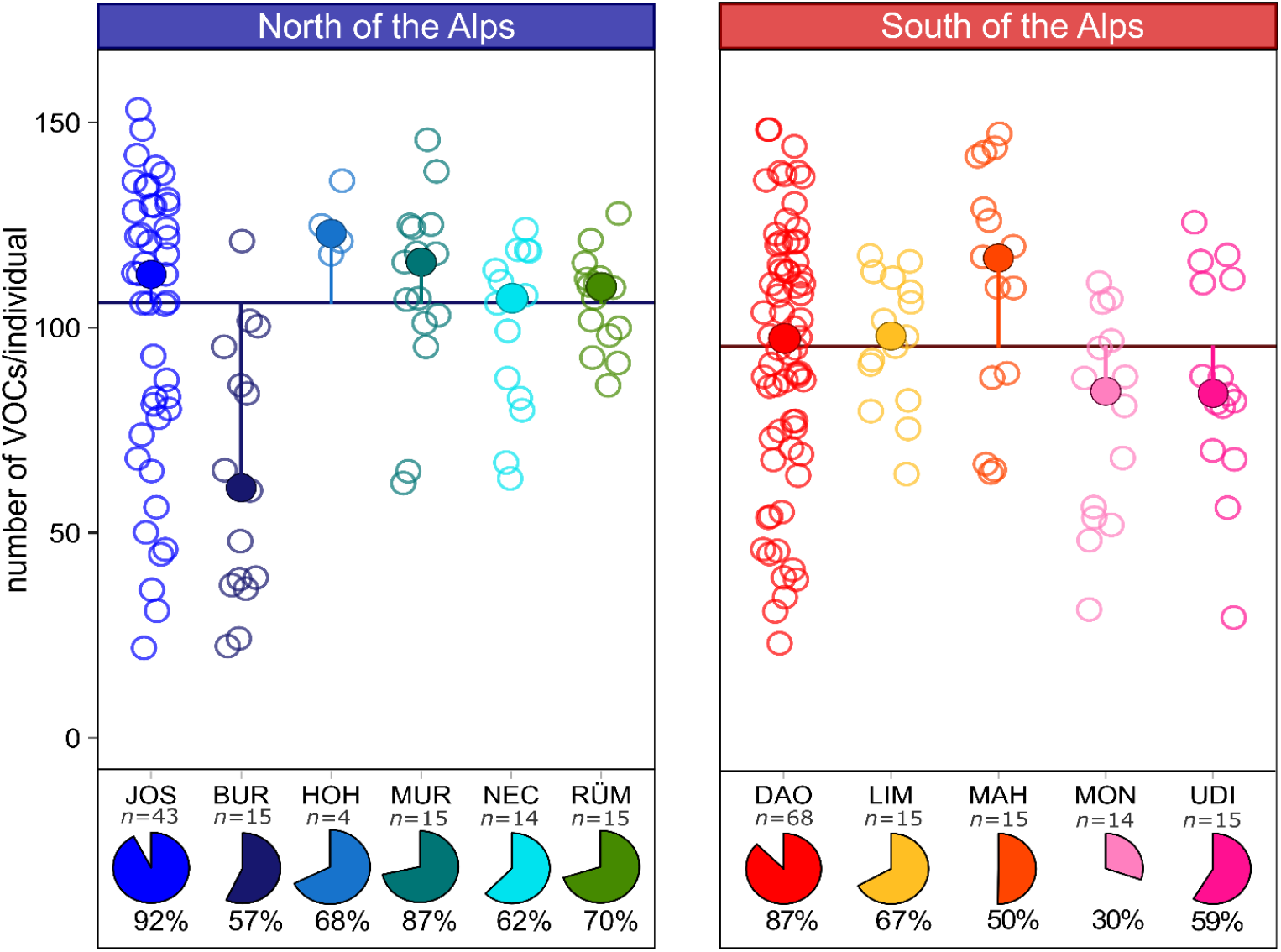
Number of floral scent compounds recorded in *Arum maculatum* individuals from populations north and south of the Alps, respectively. Filled circles denote the population median of number of volatiles per individual; the vertical lines indicate the distance to the region median (horizontal line); and open circles mark the number of volatiles detected in the individual samples. Pie charts indicate the percentage of volatiles detected per population (*n*, sample size) compared to the number of compounds detected across all samples (291 compounds). See Fig. 1 and Table S1 for identification of population codes.

**Table 1.** Median amounts of total absolute and relative (contribution of single compounds to total scent) inflorescence scent of *Arum maculatum* surveyed in six and five populations north and south of the Alps, respectively. North and South columns (bold headers) present the regional median of the corresponding populations (following columns). Volatiles with a median amount of <1% in any population are pooled.

| KRI | Compound name | **North**<br>(n=106) | JOS<br>(n=43) | BUR<br>(n=15) | HOH<br>(n=4) | MUR<br>(n=15) | NEC<br>(n=14) | RUM<br>(n=15) | **South**<br>(n=127) | DAO<br>(n=68) | LIM<br>(n=15) | MAH<br>(n=15) | MON<br>(n=14) | UDI<br>(n=15) |
| --- | --- | --- | --- | --- | --- | --- | --- | --- | --- | --- | --- | --- | --- | --- |
|  | Median total absolute amount of scent trapped (ng inflorescence<sup>-1</sup> h<sup>-1</sup>) | 67.4 | 167.2 | 13.0 | 565.8 | 80.7 | 39.4 | 41.7 | 214.7 | 311.4 | 203.8 | 196.9 | 201.4 | 42.3 |
|  | Total number of volatiles | 283 | 269 | 166 | 197 | 204 | 181 | 208 | 265 | 254 | 195 | 146 | 88 | 171 |
|  | Aliphatic components |  |  |  |  |  |  |  |  |  |  |  |  |  |
| 893 | 2-Heptanone* | 1.4 | 1.4 | 2.4 | 0.8 | 1.3 | 1.1 | 1.4 | 6.9 | 9.3 | 0.3 | 2.9 | 11.9 | 4.0 |
| 902 | 2-Heptanol* | 0.1 | 0.1 | 0.3 | 0.3 | 0.2 | 0.2 | 0.1 | 1.2 | 1.9 | tr | 0.8 | 2.5 | 0.5 |
| 982 | 1-Octen-3-ol* | 1.9 | 2.4 | tr | 2.3 | 2.0 | 1.8 | 1.3 | 0.3 | 0.4 | tr | 6.4 | tr | 1.3 |
| 1096 | 2-Nonanone* | 0.2 | 0.1 | 0.1 | 0.1 | 0.2 | 0.1 | 0.2 | 0.7 | 0.9 | tr | 0.2 | 1.3 | 0.4 |
|  | 23 more aliphatic components <1% | 0.5 | 0.5 | 0.4 | 2.0 | 1.7 | 1.3 | 0.5 | 0.7 | 0.9 | 0.9 | 2.0 | 1.0 | 0.7 |
|  | Aromatic components |  |  |  |  |  |  |  |  |  |  |  |  |  |
| 1076 | *p*-Cresol* | 4.2 | 1.8 | 0.1 | 19.4 | 11.9 | 9.2 | 1.5 | 0.5 | 0.5 | 0.3 | 0.7 | 0.6 | 0.9 |
|  | 4 more aromatic components <1% | tr | tr | tr | 0.6 | 0.3 | 0.1 | 0.1 | tr | tr | tr | 0.1 | tr | tr |
|  | C5-branched chain components |  |  |  |  |  |  |  |  |  |  |  |  |  |
|  | 4 C5-branched chain components <1% | tr | tr | tr | 0.1 | tr | 0.2 | 0.1 | tr | tr | tr | tr | tr | tr |
|  | Nitrogen-bearing components |  |  |  |  |  |  |  |  |  |  |  |  |  |
| 965 | β-Lutidine | 0.2 | 0.1 | 0.7 | 0.2 | 0.4 | 0.4 | 0.3 | 0.1 | tr | 0.6 | 0.1 | tr | 1.3 |
| 1310 | Indole* | 24.2 | 22.3 | 20.8 | 12.6 | 24.6 | 33.4 | 35.6 | 11.9 | 11.9 | 24.8 | 8.8 | 12.3 | 9.5 |
|  | 5 more nitrogen-bearing components <1% | 0.1 | 0.1 | tr | 0.1 | tr | tr | 0.3 | 0.1 | tr | 0.1 | 0.1 | tr | tr |
|  | Irregular terpene |  |  |  |  |  |  |  |  |  |  |  |  |  |
|  | 3 irregular terpenes <1% | tr | tr | 0.4 | 0.3 | tr | 0.6 | 0.1 | 0.1 | 0.1 | 0.2 | tr | 0.3 | 0.1 |
|  | Monoterpenoids |  |  |  |  |  |  |  |  |  |  |  |  |  |
| 914 | 3,7-Dimethyloct-1-ene* | 1.5 | 1.2 | 1.1 | 2.6 | 2.1 | 1.8 | 1.7 | 4.0 | 4.3 | 4.1 | 2.4 | 4.6 | 2.7 |
| 935 | α-Citronellene*§ | 0.4 | 0.3 | 0.5 | 0.9 | 0.5 | 0.5 | 0.4 | 1.3 | 1.3 | 1.9 | 1.1 | 1.3 | 1.0 |
| 949 | β-Citronellene*§ | 4.2 | 3.5 | 8.0 | 11.6 | 3.4 | 7.1 | 3.5 | 9.7 | 10.8 | 10.1 | 6.5 | 9.9 | 8.2 |
| 972 | 3,7-Dimethylocta-2-ene* | 3.1 | 1.8 | 2.8 | 5.1 | 9.6 | 2.6 | 3.3 | 4.3 | 4.5 | 4.4 | 5.6 | 2.1 | 3.2 |
| 982 | Sabinene* | 0.2 | tr | 1.0 | 0.2 | tr | 0.4 | 0.4 | 0.4 | 0.3 | 1.4 | 0.3 | 1.2 | tr |
| 1005 | 2,6-Dimethylocta-2,6-diene* | 1.2 | 0.9 | 1.0 | 3.4 | 4.1 | 1.0 | 1.5 | 1.7 | 1.8 | 1.7 | 1.9 | 0.5 | 1.6 |
| 1076 | Dihydromyrcenol | tr | tr | tr | tr | tr | tr | tr | 0.4 | 0.4 | 1.0 | tr | 0.2 | 0.5 |
|  | 21 more monoterpenoids <1% | 0.3 | 0.4 | tr | 2.2 | 0.9 | 0.6 | 0.9 | 0.6 | 0.7 | 1.9 | 1.2 | 0.4 | 1.1 |
| Sesquiterpenoids |  |  |  |  |  |  |  |  |  |  |  |  |  |  |
| 1357 | Bicycloelemene | 0.4 | 0.5 | 0.1 | 0.5 | 1.9 | 0.1 | 0.5 | 0.2 | 0.1 | 0.2 | 0.6 | 0.1 | 0.9 |
| 1399 | α-Copaene* | 1.0 | 1.8 | 1.3 | 0.5 | 0.5 | 0.7 | 0.8 | 0.8 | 0.6 | 1.0 | 1.0 | 1.6 | 0.6 |
| 1434 | Isocaryophyllene | 0.9 | 1.3 | 0.7 | 0.4 | 0.6 | 0.5 | 1.1 | 0.9 | 0.7 | 1.2 | 1.3 | 1.2 | 0.8 |
| 1450 | β-Caryophyllene* | 3.0 | 5.5 | 3.0 | 2.3 | 1.4 | 2.7 | 2.7 | 2.9 | 2.2 | 3.0 | 4.0 | 5.3 | 2.8 |
| 1484 | α-Humulene* | 2.8 | 4.7 | 2.8 | 1.7 | 1.2 | 2.3 | 2.7 | 2.3 | 1.6 | 2.5 | 3.0 | 4.0 | 2.6 |
| 1501 | Germacrene D* | 0.9 | 1.3 | 1.4 | 0.3 | 0.3 | 0.9 | 0.7 | 0.5 | 0.3 | 0.5 | 0.7 | 1.3 | 0.7 |
| 1520 | Bicyclogermacrene | 0.9 | 1.0 | tr | 0.6 | 2.1 | tr | 1.3 | 0.4 | 0.2 | 0.2 | 1.3 | tr | 1.7 |
| 1547 | δ-Cadinene | 1.2 | 1.9 | 1.4 | 0.6 | 0.6 | 1.5 | 1.1 | 0.4 | 0.3 | 0.5 | 0.7 | 0.4 | 1.0 |
|  | 10 more sesquiterpenoids <1% | 0.4 | 0.5 | 0.3 | 0.4 | 0.3 | 0.5 | 0.6 | 0.3 | 0.1 | 0.3 | 0.4 | 0.7 | 0.3 |
| Unknown compounds |  |  |  |  |  |  |  |  |  |  |  |  |  |  |
| 829 | UNK 829 *m/z*: 54,67,110,41,81,39 | 0.3 | 0.8 | tr | 0.2 | 0.2 | 0.3 | 0.1 | tr | tr | tr | 2.0 | tr | 0.3 |
| 1394 | UNK 1394 *m/z*: 69,55,41,82,95 | 0.2 | 0.2 | tr | 0.1 | 0.6 | 0.2 | 0.2 | 0.1 | 0.1 | 0.1 | 1.1 | 0.2 | 0.1 |
| 1409 | UNK 1409_1 *m/z*: 81,55,67,95,41 | 0.2 | 0.2 | tr | 0.4 | 0.4 | 0.1 | 0.1 | 0.2 | 0.2 | 0.1 | 0.6 | 0.2 | 1.1 |
| 1415 | UNK 1415 *m/z*: 69, 81,41,95,55 | 3.7 | 3.9 | 1.7 | 2.3 | 7.3 | 3.7 | 3.4 | 3.8 | 3.1 | 2.8 | 10.4 | 2.3 | 11.3 |
| 1492 | UNK 1492 *m/z*: 105,161,91,41,93 | 1.7 | 2.7 | 1.4 | 0.3 | 0.5 | 0.5 | 1.8 | 1.2 | 1.0 | 1.4 | 1.6 | 3.1 | 0.6 |
| 1503 | UNK 1503 *m/z*: 81,107,163 | 0.8 | 0.9 | 0.2 | 0.4 | 0.8 | 0.6 | 1.0 | 0.2 | 0.2 | 0.1 | 0.4 | 0.2 | 0.3 |
| 1524 | UNK 1524 *m/z*: 105,161,204,119,93 | 0.7 | 1.0 | 1.3 | 0.2 | 0.2 | 0.8 | 0.7 | 0.4 | 0.2 | 0.4 | 0.5 | 0.9 | 0.5 |
| 1699 | UNK 1699 *m/z*: 81,163,191,95,123 | 3.6 | 4.1 | 3.0 | 0.5 | 1.8 | 3.2 | 5.2 | 1.3 | 1.1 | 1.6 | 2.0 | 0.8 | 3.2 |
|  | 192 more unknowns <1% | 2.9 | 3.9 | 1.6 | 4.7 | 4.9 | 3.8 | 4.9 | 2.7 | 2.2 | 4.8 | 4.6 | 3.4 | 3.7 |

Volatiles are ordered according to compound class, and within class by Kováts' retention index (KRI). The total number of volatiles is also given.
Abbreviations: \* = identification of compound was verified by authentic standards; tr = trace relative amount (<0.05%); *m/z* = mass-to-charge ratio in decreasing order of abundance. North: JOS, Josefiu; BUR, Burg Hohenstein; HOH, Hohendilching; MUR, Murnau; NEC, Horb am Neckar; RÜM, Rümikon; South: DAO, Daone; LIM, Limone-Piemonte; MAH, Santa Maria Hoè; MON, Montese; UDI, Udine.
§ Synthetic (+)- $\alpha$ - and (+)- $\beta$ -Citronellene coeluted with natural detected $\alpha$ - and $\beta$ -Citronellene on a chiral column (MEGA-DEX DMT Beta SE, 30m × 0.25mm ID, 0.23 $\mu$ m film) (Gfrerer *et al.* unpublished).

The absolute amounts of single compounds significantly differed both between regions (permANOVA: pseudo-*F*_1,222_ = 22.52, *P* < 0.001), JOS and DAO only (pseudo-*F*_1,109_ = 9.96, *P* < 0.001), and among populations within regions (pseudo-*F*_9,222_ = 6.44, *P* < 0.001), but differences were more pronounced between regions than among populations within regions (*north* vs. *south* OOB error: 10.3%; among populations within *north* OOB error: 28.3%; within *south* OOB error: 23.6%). Only a few abundant compounds dominated the scent bouquet of *A. maculatum*, including indole, *β*-citronellene, the unknown UNK1415, and 3,7-dimethyloct-2-ene (all abundant in both regions), *p*-cresol (most abundant only north), and 2-heptanone (only south, Table **1**).

We also detected differences in the relative amounts of scent compounds between regions (permANOVA: pseudo-*F*_1,222_ = 30.18, *P* < 0.001), JOS and DAO only (pseudo-*F*_1,109_ = 22.79, *P* < 0.001), and among populations within regions (pseudo-*F*_9,222_ = 4.90, *P <* 0.001; Fig. **3**); but, again, these differences were more pronounced at the inter-regional than within-region levels (*north* vs. *south* OOB error: 9.0%; among populations within *north* OOB error: 33.9%; among populations within *south* OOB error: 25.2%; see also Supporting Information Figure **S1**).

Across all populations, variation in absolute or relative amounts of scent could not be explained by their geographic distances (Mantel’s *Rho* = 0.108, *P* = 0.25 and *Rho =* −0.154, *P* = 0.85, respectively).

Of the 25 compounds each that were most responsible for regional differences in the absolute and relative datasets in the r*andomForest* analyses, 20 were common to both datasets (Supporting Information Table **S3**). These 20 included 2-heptanone, 2-heptanol, and *α*- and *β*-citronellene, all of which were more abundant (in relative and absolute amounts) south of the Alps, and 1-pentadecanol, the unknown UNK1503, *p*-cresol, and indole, which occurred in higher amounts north of the Alps (Table **1**, and Supporting Information Tables **S2, S3**). Many of these compounds, but also some non-overlapping ones (absolute: *α*-copaene, *β*-caryophyllene; relative: UNK1409, bicyclogermacrene), explained most of the scent variation in scent among all samples (for relative data see Fig. **3**; for absolute data see Supporting Information Table **S4**).

**Fig 3.**
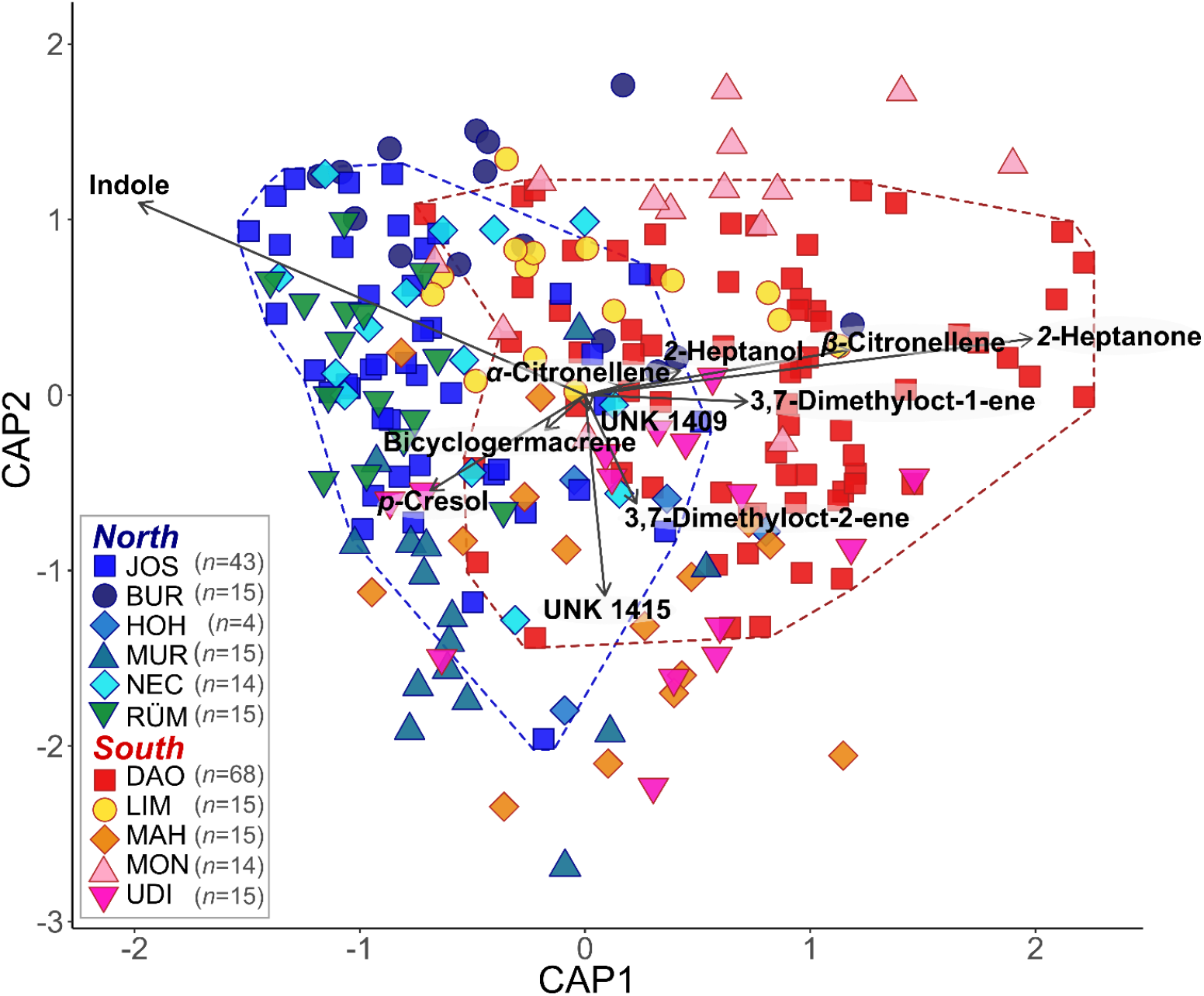
Canonical analysis of principal coordinates (CAP) based on a Bray–Curtis dissimilarity matrix of relative floral scent in *Arum maculatum* individuals from populations north and south of the Alps, respectively. *N* denotes the sample size per population. The vectors depict the volatiles most correlating with the *capscale* scores. The colored dashed lines delineate the individual scent variation of the two most extensive sampled populations JOS (blue) and DAO (red). See Fig. 1 and Table S1 for identification of population codes.

We also observed considerably high variation in scent within populations, most prominently in the most extensively sampled northern (JOS) and southern (DAO) populations, which harboured almost all of the absolute and relative scent variation of their respective region (for relative data see Fig. **3**).

### Fruit set

Of the 233 individuals surveyed for inflorescence scent, 113 set fruit in summer. Percentages of fruit set were significantly higher north of the Alps (42 ± 41% mean ± sd, 0–100% Min– Max) than south of the Alps (26 ± 33% mean ± sd, 0–100% Min–Max; Fig. **4**; *region*: *F*_1,209_ =10.11, *P* = 0.002), and differed significantly among populations within regions (*population* nested within *region*: *F*_8,209_ = 2.23, *P* = 0.03).

**Fig 4.**
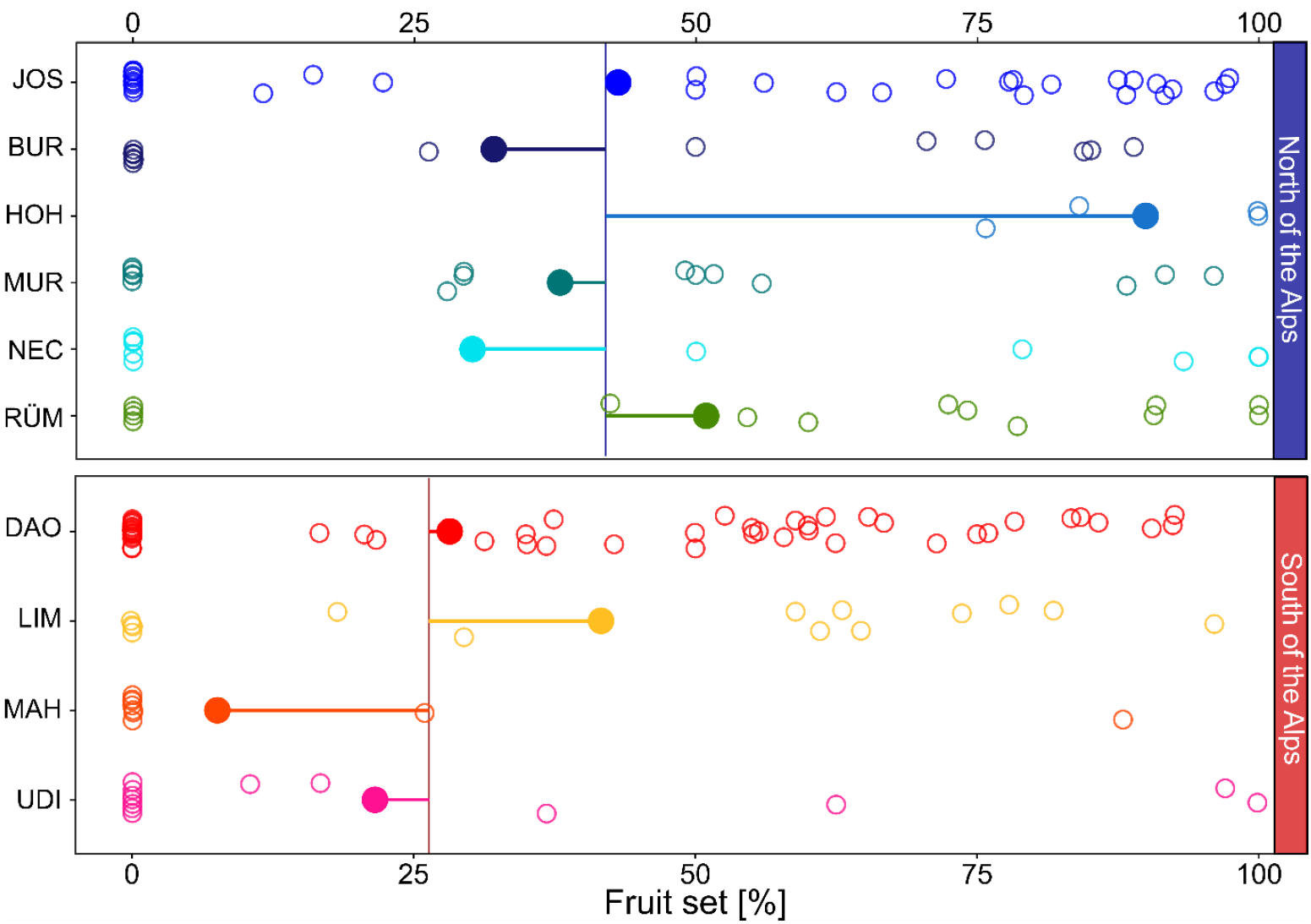
Fruit set (% female flowers that developed into fruits) of *Arum maculatum* individuals from populations north and south of the Alps, respectively. Filled circles denote the population mean of fruit set; horizontal lines indicate the distance to the region mean (vertical line); and the open circles mark the fruit set of each individual. See Fig. 1 and Table S1 for identification of population codes.

### Phenotypic selection on scent

In the most extensively sampled northern (JOS) and southern (DAO) populations, we tested 19 and three compounds for phenotypic selection, respectively, which correlated with relative fruit set in the elastic net and *Boruta* analyses (see Material and Methods; Supporting Information Methods **S3**). Of those 22 compounds, seven showed signals of linear phenotypic selection (two of which as an interaction), all in the north, and two for nonlinear (quadratic) phenotypic selection, all in the south (Fig. **5**). Seven of the overall nine compounds that were under phenotypic selection correlated positively with relative fruit set (linear: 2-heptanol, 2-nonanol, *α*-terpinene, UNK681, and UNK1496 together with UNK1503; nonlinear: sabinene), and two negatively (linear: UNK960; nonlinear: 4-terpinenol; Fig. **5** and Supporting Information Table **S5**, and Notes **S1**).

**Fig 5.**
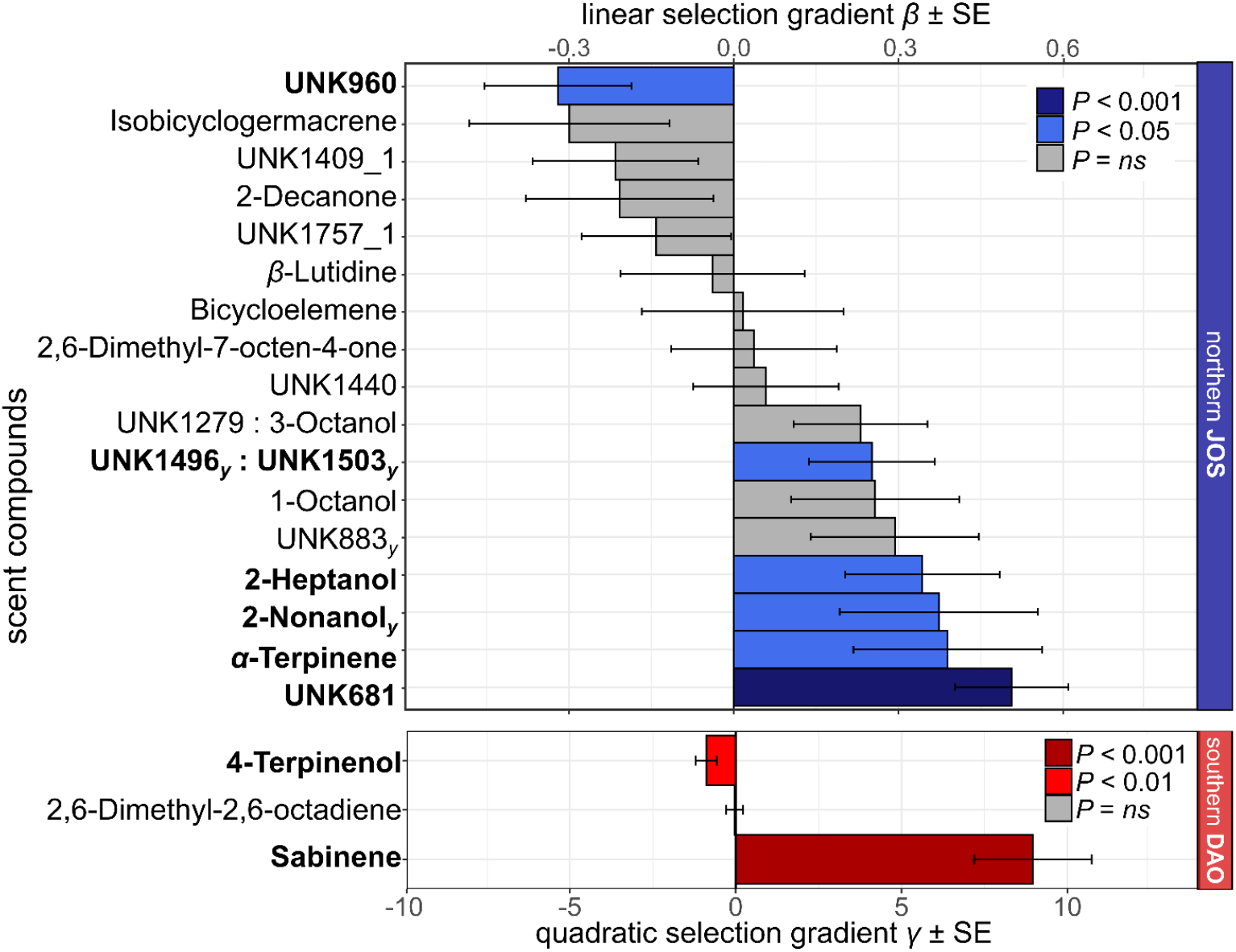
Linear selection gradients *β* and nonlinear quadratic selection gradients *γ* (and their standard errors, SE) for individual floral scent compounds in the most extensively sampled *Arum maculatum* populations from north (JOS, blue, *n* = 43) and south (DAO, red, *n* = 68) of the Alps, respectively. Only compounds that correlated with relative fruit in the elastic net/*Boruta* analyses are shown (see Material and Methods). Scent compounds under significant selection (*P* < 0.05) are in bold and their bars are coloured. Note the different scaling for linear (*β*) and nonlinear (*γ*) selection. For the northern population, compounds that were also detected by the nonlinear *Boruta* analyses are indicated with a subscript (*γ*).

Of the 25 compounds that most strongly contributed to the absolute (and relative) differences in scent between the regions each, only four were under selection (north: 2-heptanol, 2-nonanol, UNK681, UNK1503), but not others (*e*.*g*., 2-heptanone, *α*- and *β*-citronellene; Supporting Information Table **S3**). Differences in absolute and relative scent traits between the northern JOS and the southern DAO remained significant, regardless of performing PERManova analyses separately on the nine compounds that correlated with relative fruit set in the elastic net/*Boruta* and were under selection (absolute vs. relative datasets: pseudo-*F*_1,109_= 18.4 vs. 30.8, both *P* < 0.001), or on the 93 compounds that were not under selection and did not correlate with fruit set (absolute vs. relative datasets: pseudo-*F*_1,109_ = 9.8 vs. 24.5, both *P* < 0.001; see Supporting Information Fig. **S2**, Methods **S3**).

## DISCUSSION

Our study shows that *Arum maculatum* has hyperdiverse inflorescence scents that differ in their composition between populations north vs. south of the Alps. Contrary to our expectations, scent was found to differ not only among southern but also among northern populations. As expected, individuals from the southern populations had lower fruit set than northern ones, and different signs of phenotypic selection were found in the most extensively sampled northern and southern populations.

### Hyperdiversity of floral scent

With 291 floral volatiles recorded, the inflorescence scent diversity of *A. maculatum* is extraordinarily high and, to the best of our knowledge, not matched by any other plant species. In fact, we are not aware of any species from which more than 200 floral compounds are reported, a number that a single *A. maculatum* individual can reach by three quarters (max. = 152 VOCs; Fig. **2**). This difference in the number of scent compounds between *A. maculatum* and other species cannot just be explained by differences in techniques used for scent analyses, given that scents of a high number of species were analysed using a similar approach as we did (dynamic headspace and thermal desorption of samples)^60–64^. Species closest to the high number of VOCs in *A. maculatum* include the sapromyiophilous *Sauromatum guttatum* (Araceae, with altogether 196 different VOCs^65,66^), as well as the insect-pollinated and rewarding *Geonoma macrostachys* (Arecaceae, 176 VOCs^60^) and *Echinopsis ancistrophora* (Cactaceae, 145 VOCs^67^). Other species for which *c*. 100 VOCs are described likewise include insect-pollinated and rewarding species (*e*.*g*., *Acleisanthes wrightii*, Nyctaginaceae^61^; *Saraca asoca*, Fabaceae^64^; *Philodendron bipinnatifidum*, Araceae^62^; *Pyrus communis*, Rosaceae^63^), but also the sexually deceptive orchid *Ophrys sphegodes*^68^. Thus, high numbers of compounds are found across a wide range of plant families and are apparently not restricted to a specific pollination system.

One explanation for the high diversity of scent compounds in *A. maculatum* is that this species likely imitates its moth fly pollinators’ various breeding substrates, all potentially differently scented. The two main pollinators, *Psychoda phalaenoides* and *P. grisescens*, breed in a variety of different substrates such as rotting manure from cattle and horse, fungi (*P. grisescens*), waste pits, mud-flats, plant litter in drainages and ditches (*P. phalaenoides*), and in the hygropetric zones of riverbanks and ponds^69–76^. *Arum maculatum* emits compounds described from quite a number of such substrates, *e*.*g*., cattle and horse manure (*e*.*g*., indole, *p*-cresol, skatole), fungi (1-octen-3-ol, (*E*)-2-octen-1-ol, 3-octanone), and general degrading and fermenting plant or animal material (*e*.*g*., 2,3-heptanedione, acetoin, butanoic acid)^40,77–79^. Highly specialised deceptive plant systems frequently rely on only a few volatiles to attract pollinators – they seem to imitate a more specific model, and thus, release less complex scent blends^*e*.*g*., 40,80–83^.

The number of volatiles detected across the 233 individuals (11 populations) of *A. maculatum* (291 VOCs) is five to ten times higher than previously reported for this species (18– 61, and 143 VOCs in total^53–58,84^). This discrepancy cannot be explained by differences in sample size, as a similar number of individuals were surveyed in those previous studies (*n* = 222 in total, representing 23 populations). Interestingly, we found a similar number of compounds in some individuals (up to 152 VOCs; median of 102; Fig. **2**) as overall detected previously^53–58,84^. With the exception of two studies^56,58^, each sharing one of our sampled populations (JOS and MON, respectively), all previous studies sampled scents in other populations across Europe^53–55,57^. Thus, some of the differences in the number of *A. maculatum* compounds detected across studies might reflect population-specific scent characteristics (see Fig. **2**). More importantly, however, we believe that the discrepancy in the number of compounds recorded largely reflects differences in methodology between the present and previous studies. These are, for example, higher sensitivity of modern GC/MS systems; usage of more selective adsorbent agents (Carbotrap/Tenax-TA vs. solid-phase micro-extraction^55,57^ vs. Twister^58^); *in situ* vs. *ex situ* sampling^56,84^; and including all vs. only compounds above a specific threshold in relative amounts^55,58^. Of the 92 compounds chemically identified in this study, more than half (50) were previously unknown to be released by *A. maculatum*. Some of these newly described compounds for *A. maculatum* are known from other species of Araceae (*e*.*g*., nerol, (*E,E*)-*α*-farnesene, 6-methyl-5-heptene-2-ol, γ*-*terpinene; *Sauromatum guttatum*^65^, *Anthurium* spp.^85,86^) or other plant families (*e*.*g*., methyl anthranilate, isobutyl butyrate, citronellal^4,6^). However, this study is, to the best of our knowledge, the first to identify *p*-cresyl butyrate as a floral scent compound.

### Geographic patterns of floral scent

The qualitative, absolute, and relative differences in scent among populations of *A. maculatum* from south of the Alps may be explained by the fact that the pollinator assemblages there are more diverse in terms of abundance, species composition, and sex ratio^52^, ^Laina, D. *et al*., unpubl.^. North of the Alps, females of *P. phalaenoides* are the principal pollinators in all studied populations^52^, ^Laina, D. *et al*., unpubl.^, even though other *Psychoda sp*. also occur in this region^55,71, Laina, D. *et al*., unpubl^; hence, the scent variation we observed among northern populations is not reflected by variations in pollinator spectra in this region. In the present study, all variations in scent were more pronounced between regions than among populations within each region. This strong regional component of scent variation in *A. maculatum* across the Alps thus accords with strong differences in pollinator spectra^52, Laina, D. *et al*., unpubl.^, but also coincides with a weak genetic (AFLP) subdivision of *A. maculatum* across this geographic barrier^59^. Previous studies in *A. maculatum* also found population effects in scent composition^55,57,58^; however, our study is the first to demonstrate such population differentiation in scent across the Alps. Intraspecific variation in floral scent among populations and regions has also been reported for other plant species^14,15,34,37,67^. In some of those, this variation, as in our study, could be linked to pollinator assemblages and/or genetic patterns^15,34,37^, but not in others^14,67^.

### Phenotypic selection on floral scents

The two most extensively sampled northern (JOS) and southern (DAO) populations differed in absolute and relative amounts of scent, regardless of whether the analyses were conducted on all compounds, only on those that correlated with relative fruit set and were under selection, or those that did not correlate with fruit set (Material and Methods, Supporting Information Fig. **S2**). Thus, this regional difference in scent could be caused by different selection regimes, as well as other reasons, such as phenotypic plasticity (but see^58^) or genetic drift^87,88^. In support of differential selection, we detected population-specific signatures of phenotypic selection on scent in JOS and DAO, possibly due to different olfactory preferences of those *Psychoda* species that dominate the pollinator spectra of *A. maculatum* in the northern (female *P. phalaenoides*) vs. southern (and *P. grisescens*) regions^52,55,58,Laina, D. *et al*., unpubl.^.

For the five compounds under phenotypic selection that we were able to chemically identify (*i*.*e*., 2-nonanol, 2-heptanol, sabinene, 4-terpinenol and *α*-terpinene), information on their attractiveness to pollinators of *A. maculatum* is lacking. However, the aliphatic compounds 2-heptanol and 2-nonanol are known, together or alone, as attractants for bees (Meliponini^89,90^) and kleptoparasitic flies^91^, and as (sex-)pheromones of female Diptera (Cecidomyiidae^92^) and female non-Diptera (Trichoptera^93^). The monoterpenoids sabinene, *α*-terpinene, and 4-terpinenol are defence substances of some insects (Coleoptera^94,95^, Lepidoptera^96^) that repel Coleoptera^97^, but are used by Hymenoptera^98^ and Lepidoptera^99^ for host-finding and as oviposition stimulants. The latter two are also pheromones of fruit flies^100^. In summary, these five compounds, found to be under phenotypic selection, elicit responses in insects other than moth flies. Further, they are known as floral scent from other sapromyiophilous species^65,66,77,81,101,102^, and some of them (*α*-terpinene and 4-terpinenol) are also known from cattle dung^79,103^, *i*.*e*., one of the oviposition substrates of moth flies. Further research is required to establish whether these five compounds, which are all widespread floral scent compounds^4,6^, are attractive to the pollinators of *A. maculatum*.

Several of the compounds most responsible for regional differences in inflorescence scent (*e*.*g*., 2-heptanone, 3,7-dimethyloct-1-ene, UNK966; Supporting Information Table **S3**) did not show signals of phenotypic selection (Fig. **5**). Thus, the different selection regimes cannot explain several of the obvious differences in scent between *A. maculatum* from north and south of the Alps (see also Supporting Information Fig. **S1**). Some other compounds, however, which also differed in their absolute amounts between regions (2-heptanol, 2-nonanol, UNK681, sabinene; Supporting Information Table **S3, S5**) were under phenotypic selection, either in northern JOS or southern DAO (Fig. **5**), and some of the differences between regions could therefore be due to differential selection.

Somewhat unexpectedly, we did not find phenotypic selection for the most abundant compounds in the scent of *A. maculatum* (*e*.*g*., indole, *β*-citronellene, unknown UNK1415), with the exception of 2-heptanol (Fig. **5**). Even more surprisingly, we also did not find phenotypic selection for those compounds known to attract *P. phalaenoides* (*i*.*e*., indole, 2-heptanone, *p*-cresol, *α*-humulene^50,84^), occurring both north and south of the Alps, and also in JOS and DAO^52^. This is in contrast to most other studies, where main compounds and/or pollinator attractants^29,32,33,35^ showed signals of phenotypic selection (but see^31,34^). Possible explanations for not finding phenotypic selection on the main compounds of *A. maculatum* include: (1) their release in amounts high enough to achieve maximum pollinator attractiveness (see also^34^); (2) opposing selection pressures on these compounds by different pollinators or herbivores, resulting in zero ‘net’ selection^33,34^; or that (3) their relationship with flower visitors is nonlinear and nonquadratic^104–106^. Although our multivariate models detected nonlinear phenotypic selection by including quadratic terms, such quadratic analyses cannot uncover all potential nonlinear relationships (*e*.*g*.,^107^). Hence, we cannot exclude the possibility that such abundant and/or attractive compounds are still under phenotypic selection, which in turn calls for future statistical developments that allow testing for any kind of nonlinear multivariate relationships.

Deceptive plant species might experience stronger selection than rewarding ones^47^. However, by comparison with rewarding species^30–36^, we found that deceptive *A. maculatum* does not release a higher number of volatiles with signatures of phenotypic selection (7% vs. 3–42%), but they appear to be under slightly stronger positive linear phenotypic selection (−0.3−0.5 vs. −0.3−0.4^30–35^ Min−Max) and stronger nonlinear phenotypic selection (−0.9−9.0 vs. −0.5−-0.3^36^ Min−Max; Fig. **5**). Future studies on other deceptive plant species that also attract specific pollinators by chemical cues but have lower levels of fruit set than *A. maculatum* (such as many orchids^44,45^) might reveal even stronger signatures of phenotypic selection.

## Conclusions

Our study on sapromyiophilous *Arum maculatum* reports the highest number of floral volatiles ever found in a single plant species to date. This chemical hyperdiversity could be due to the fact that this brood-site deceptive plant species imitates the odours of a multitude of differently scented breeding substrates of its moth fly pollinators (*e*.*g*., dung, fungi, rotting plant material). We recorded pronounced scent differences between populations from north vs. south of the Alps, and this geographic pattern in scent agrees with previously described pollinator and genetic patterns across this geographic barrier. Our results provide, for the first time, evidence that floral scents of a deceptive plant are under phenotypic selection and suggest that populational/regional differences in scent are partly due to differential selection, while other reasons, such as phenotypic plasticity and genetic drift cannot be excluded. In *A. maculatum* and other plants where phenotypic selection on scent was demonstrated^29–36^, the biological role of most compounds under selection is unknown and awaits determination in future studies.

## MATERIALS AND METHODS

### Study species and populations

Brood-site deceptive *Arum maculatum* L. (Araceae) is a rhizomatous perennial woodland herb (2*n* = 4x = 56) that is widespread throughout Western and Central Europe, including the British Isles, and reaches as far south as Italy, Northern Spain, and the Balkans^52,108,109^. It exhibits a sapromyiophilous pollination strategy, is thermogenically active, and emits a strong dung-like scent for attracting moth fly pollinators during the evening on the first day of anthesis^53,56,110,111^. The inflorescence of *Arum maculatum* consists of a spadix (fleshy spike) and a spathe (bract), is protogynous, and the anthesis lasts less than two days^51,56,110^. The spathe, which completely encloses the spadix during floral development, partially opens during anthesis to reveal the sterile appendix of the apical part of the spadix. This appendix produces and releases the scent for pollinator attraction^50,51,53,84^. At the base of the spadix, female (fertile and sterile) flowers are situated lowest, followed upwards by male flowers and staminodes (sterile male flowers). All flowers remain enveloped by the spathe during anthesis, forming a chamber that is closed by the staminodes throughout the female stage to prevent trapped insects from leaving. Pollinators are attracted in the evening on the first day of anthesis, during the female stage, slip and fall into the floral chamber, and are trapped overnight^51,52,55,111^. On the next morning, during the male stage, they are dusted with pollen, before being released at around noon when the staminodes and spathe wither^51,52,110^. After pollination in spring, red berry-like fruits develop as an infructescence until summer^112^.

In 2017–2019, during springtime, we collected scent from randomly chosen *A. maculatum* individuals of six populations located north of the Alps (*n* = 106; Northwestern Austria: JOS; Central/Southern Germany: BUR, HOH, MUR, NEC; Northern Switzerland: RÜM) and five populations from south of the Alps (*n* = 127; Northern Italy: DAO, LIM, MAH, MON, UDI) (Fig. **1**). We kept a minimum distance of one metre between sampled individuals to avoid sampling potential clones, as *A. maculatum* can propagate vegetatively by fragmenting rhizomes^51^. In summer, we harvested fruits from all individuals surveyed for scent. At most sites, we recorded fruit set of 15 individuals, except for each of the largest population per region (JOS and DAO; *n* = 70 each), and a northern population (HOH; *n* = 7) where only a few individuals had flowered at the time of scent sampling (Fig. **1** and Supporting Information Table S**1**).

### Plant volatile collection and analysis

Scent sampling took place on the first day of anthesis during the female stage between 6 pm and 7.30 pm, the period of maximum scent emission^56^, employing a non-invasive dynamic headspace technique. We enclosed each inflorescence *in situ* using an odourless plastic oven bag (c. 30×12 cm; Toppits^®^, Melitta, Germany) and immediately collected scent for five minutes at 200 ml min^-1^ on adsorbent tubes (inner diameter: 2 mm) filled with a mixture of Tenax-TA (mesh 60–80) and Carbotrap B (mesh 20–40; 1.5 mg each; both Supelco, Germany), using a battery-operated vacuum pump (rotary vane pump G12/01 EB, Gardner Denver Austria GmbH, Vienna, Austria)^56^. In the same way, we collected scent samples from leaves and ambient air as negative controls in each population.

The dynamic headspace samples were analysed by thermal desorption-gas chromatography/mass spectrometry (TD-GC/MS)^56^, and obtained data were handled using *GCMSolution* v.4.41 (Shimadzu Corporation, Kyoto, Japan) (for details see Supporting Information Method **S1**). Compounds were chemically identified by comparison of Kováts’ retention indices (KRIs^113^), based on commercially available *n*-alkanes (C_7_–C_20_), and mass spectra to data available in the libraries of Adams^114^, FFNSC 2, Wiley9, NIST11, and ESSENTIAL OILS (available in *MassFinder 3*, Hochmuth Scientific Consulting, Hamburg, Germany). We established an own library of mass-spectral and KRIs for semi-automatic analysis (Supporting Information Method **S1**). Whenever possible, compounds were verified by comparison to authentic reference standards available in the collection of the Plant Ecology Lab of Salzburg University, or to chemically synthesised reference compounds (Supporting Information Method **S2**). Of the 267 collected scent samples, 233 yielded a sufficiently informative chromatogram and were included in the analysis (Fig. **1**). Ultimately, a compound was only considered if it occurred in more than three scent samples and did not occur in leaf and air controls.

### Fruit set

Percentage fruit set (*i*.*e*., number of fruits/total number of flowers per individual × 100) was determined as a measure of female reproductive success. For selection analyses we further estimated relative fruit set (*i*.*e*., number of fruits per individual/ mean number of fruits per given population) as a measurement of female reproductive success^32,35^, standardised per population for the most extensively sampled populations JOS and DAO. In one southern population (MON), a shallow landslide destroyed all plants, with the exception of one; hence, this population was excluded from fruit set analyses.

### Statistical analyses

#### Geographic patterns in scent and fruit set data

In order to test for geographic differences in floral scent, we performed permutational multivariate analyses of variance (permANOVAs^115^) as implemented in the R package *vegan* v.2.6-6^116^. We did this on (1) pairwise Bray–Curtis dissimilarities of either absolute or relative scent data (*i*.*e*., absolute amount of single compounds or relative amount of single compounds in relation to the total amount of scent in a sample, respectively); and (2) Euclidean distances of both total absolute emission of scent and of total number of floral volatiles per individual. In all these analyses, we used *region* (north vs. south of the Alps) and *population* nested in *region* as explanatory variables (9,999 permutations). Using permANOVA (*population* as explanatory variable, 9,999 permutations), we also tested for differences in relative and absolute scent, and for geographic patterns of selection in the two most extensively sampled northern (JOS) and southern (DAO) populations. To this aim, we either used all compounds, or only those that were under selection and correlated with relative fruit in the *elastic net/Boruta* (see below and Supporting Information Methods **S3**), or those that were not under selection and did not correlate with relative fruit set.

The Bray–Curtis dissimilarity matrices (based on absolute and relative scent data across all populations) were further used to conduct constrained analyses of principal coordinates (CAP^117^) with *population* as factor, using the *capscale* function in *vegan* to visualize similarities and dissimilarities in scent among the samples (following^118,119^). For each ordination, we also calculated vectors, representing compounds most correlating with the axes (Pearson correlations with *capscale scores, r* > |0.5|, corrected for false-discovery rate^120^). Given that CAP is not appropriate to display similarities and dissimilarities in scent between only two populations in a two-dimensional ordination, we used non-metric multidimensional scaling (nDMS) to visualize similarities and dissimilarities in scent among the samples of only JOS and DAO, using only compounds that correlated with relative fruit set or those that did not.

Additionally, we subjected the absolute and relative scent data to random forest analyses^121^ by the R package *randomForest* v.4.6-14^122^ (*ntree* = 9,999 bootstrap samples with *mtry* = 17) to evaluate the distinctness in scent of northern and southern samples (factor *region*) and among populations within each region (factor *population*)^123^. Distinctness was quantified as the average out-of-bag (OOB) error estimate (in %), *i*.*e*., the more distinct, the lower the OOB error. From the resulting *randomForest* objects, we further extracted the *importance* measurements to determine volatiles that are critical for regional distinction.

To test for relationships between the dissimilarity of median absolute and relative scent properties of populations and their geographic distances (in km), we performed Mantel tests with the function *mantel* in *vegan* (9,999 permutations, Spearman’s rank correlation). To assess whether absolute amounts of single compounds under selection (see below) differ between the two regions, we performed Mann–Whitney U tests. Differences in fruit set across regions and among populations within regions were assessed by a generalised linear model (*regions* and *populations* nested within *regions* as factors) that was analysed by an ANOVA.

#### Analyses of phenotypic selection

To estimate the direction and strength of phenotypic selection on scent compounds, we tested for phenotypic selection^124^ in the northern JOS and southern DAO by correlating relative fruit set with z-transformed scent data (standardised to mean = 0, sd=1)^30–32,35^. These two populations cover a large part of their respective corresponding regional scent variation (see Fig. **3**). As a major challenge, our dataset has a considerable higher number of factors (VOCs) than samples. Previous studies solved this by pre-selecting variables to reduce high dimensionality^30,31,34^, and performed selection analyses only on the most abundant compounds^29,33^, on principal component scores^29,32,35^, or on physiologically active volatiles^34^. Because we only have very limited knowledge of attractive compounds in our study system^50,84^, the assumptions for principal component analysis are violated, and as also minor volatiles can be under selection^34^, these solutions were not suitable for our dataset. Instead, we pre-selected volatiles that correlate with relative fruit set via elastic net, *i*.*e*., a penalised multivariate linear regression^125^, and via the *Boruta*^126^ algorithm, to identify linear (elastic net) and nonlinear (*Boruta*) relationships between total absolute emission as well as the absolute emission of individual volatiles and relative fruit set (for details see Supporting Information Methods **S3**). Additionally, the scent matrix contained many zeros (non-detects), as many compounds were quite rare (*c*. 70% of VOCs in < 50% of samples). This zero-inflation can cause severe problems when fitting linear models, as estimates will be biased^127^. That is, the influence of an individual scent compound on fruit set can be either over- or underestimated, leading to potentially wrong conclusions. To quantify the impact of non-detects on elastic net estimates, we performed, before the pre-selective analyses, a simulation study for JOS and DAO separately (see Supporting Information Methods **S3**). Based on the simulation results, we obtained 93 and 81 scent compounds for JOS and DAO, respectively, each of which were then included in both the elastic net regression and the *Boruta* analyses (Supporting Information Methods **S3**). For the JOS population, elastic net and *Boruta* identified 19 and four volatiles, respectively; whereby the latter were already among the linear ones (Fig. **5**). In the southern DAO population, no volatile correlated with fruit set in the elastic net, but three in the *Boruta* analysis. None of these volatiles was detected for both populations (Fig. **5**). Also, total absolute scent amount did not correlate with fruit set in any of the analyses.

To ultimately test for phenotypic selection, we subjected those variables selected by the elastic net model (Supporting Information Methods **S3**) to multivariate linear regression (linear *β*-gradients)^124^, and those volatiles identified by the *Boruta* analyses (Supporting Information Methods **S3**) to multivariate quadratic regression (nonlinear/quadratic *γ*-gradients^124^) by squaring the terms and doubling resulting estimates^107^. For the multivariate regression model of the southern (DAO) population, we excluded the plant individual ‘DAO076’, as it was determined by Cook’s distance^128^ as an outlier influencing the model (*D*_DAO076_ = 235.4). Although elastic net handles multicollinearity well, volatiles identified to correlate with fruit set might still correlate with each other (*L*_2_ penalty, see Supporting Information Methods **S3**). We therefore also tested for multicollinearity within the multivariate regression models by calculating the variance inflation factor (VIF) (R package *car* v.3.0.8^129^) for each scent compound in each model. For the northern (JOS) model, the VIF value of various compounds was high (> 5) and for the unknowns UNK1496 and UNK1503 even exceeded 10, a threshold that indicates strong multicollinearity^130^. After including these two compounds as an interaction, the VIF values of most compounds were < 5, except for 3-octanol and UNK1279 (VIF > 6). After further including the interaction of the latter two volatiles in the model, the VIF values of all volatiles were < 4. Based on this, the final northern (JOS) model had an adjusted *R*^2^ of 0.71. For the southern (DAO) model, all VIF values were < 2 (adjusted *R*^2^ = 0.26). All statistical analyses were performed in R v.4.0.2^131^.

## Acknowledgements

We would like to thank Irmgard Schäffler for methodological support, Valerie Scheurecker, Karin Moosbrugger, Bernadette Mükisch, S. Sophie Brandauer and Christopher Gorofsky for their support in the field, and Florian Schiestl for providing data on populations in Switzerland. We also thank Robert A. Raguso for discussing hyperdiverse scents, and members of the Plant Ecology group of the University of Salzburg, especially Karin Gross and Herbert Braunschmid, for constructive comments on an earlier version of the manuscript. This study was funded by a grant from FWF (Austrian Science Fund; P30175-B29) to ACH, HPC and SD (PI). MH and WT gratefully acknowledge support from the WISS 2025 project ‘IDA-lab Salzburg’ (20204-WISS/225/197-2019 and 20102-F1901166-KZP).

## Author Contributions

SD, MG, ACH and HPC designed the research; EG and DL conducted the fieldwork; RF executed the scent sample laboratory work; EG and SD built the scent library, and EG analysed all scent and fruit set data; TT identified and synthesised unknown compounds; MH, WT, RF, SD and EG discussed statistical approaches for selection analyses; MH designed and performed the simulations, EG the selection analyses; EG wrote the first draft of the manuscript and all authors contributed to the final version.

## Data Availability

The R code for the simulation and the scent data that support the findings of this study will be available in the Dryad Digital Repository and are now available on request.

**The authors declare no competing interests**.

## Additional Information

**Supporting Information** is available for this paper.

