## Supporting Information for "Floral scents of a deceptive plant are hyperdiverse and under population-specific phenotypic selection"

**The following Supporting Information is available for this article:**

**Fig S1** Relative amounts of the most abundant compounds (see Table 1) in the inflorescence scents of *Arum. maculatum,* also indicating whether or not they are under phenotypic selection.

**Fig S2** Non-metric multidimensional scaling of relative (a, b) and absolute (c, d) scent patterns of *Arum maculatum* with VOCs that correlated with relative fruit (nine VOCs; a, c) or only with those that did not (93 VOCs; b, d).

**Table S1** Locality information on the sampled populations.

**Table S2** Complete list of inflorescence scent compounds detected in *Arum maculatum* (median relative amount per population and region). ***See extra Excel File***

**Table S3** Volatiles important for regional distinction in relative and absolute scent bouquets.

**Table S4** Volatiles most correlating with the *capscale* scores in absolute scent samples

**Table S5** Selection gradients $\beta$ and $\gamma$ for volatiles of *Arum maculatum.*

**Methods S1** Scent analysis (TD-GC/MS), quantification, and set-up of scent library.

**Methods S2** Synthesis of reference samples: 2,6-dimethylocta-2,6-diene, 3,7-dimethyloct-2-ene and 2,6-dimethylocta-1,7-diene ($\alpha$-citronellene)

**Methods S3** Simulation study to quantify impact of non-detects on selection estimates, and elastic net and *Boruta* analyses to pre-select scent compounds that correlate with fruit set.

**Notes S1** Mass spectrometry of unknown volatiles with significant phenotypic selection gradients.

**Fig. S1** Pie charts showing the relative amounts (median per population) of the most abundant scent compounds (see Table 1) in the inflorescence scents of *Arum maculatum* populations north (blue) and south of the Alps (red), respectively (see Table S1 for identification of population codes). Coloured outer circle sectors indicate compounds under phenotypic selection in the most extensively sampled northern JOS or southern DAO population (orange 29–36), or under no selection in those populations (grey, 1–28). Identification of compound numbers: (1) UNK829; (2) 2-Heptanone; (3); 3,7-Dimethyloct-1-ene; (4) α-Citronellene; (5) *β*-Citronellene; (6) *β*-Lutidine; (7) 3,7-Dimethyloct-2-ene; (8) 1-Octen-3-ol; (9) 2,6-Dimethylocta-2,6-diene; (10) Dihydromyrcenol (11) *p*-Cresol; (12) 2-Nonanone; (13) Indole; (14) Bicycloelemene (15) UNK1394; (16) *α*-Copaene; (17) UNK1409_1; (18) UNK1415; (19) Isocaryophyllene; (20) *β*-Caryophyllene; (21) *α*-Humulene; (22) UNK1492; (23) Germacrene D; (24) Bicyclogermacrene; (25) UNK1524 ; (26) *δ*-Cadinene; (27) UNK1699; (28) 2-Heptanol; (29) 2-Nonanol; (30) *α*-Terpinene; (31) UNK681; (32) UNK960; (33) UNK1496; (34) UNK1503; (35) 4-Terpinenol; (36) Sabinene.

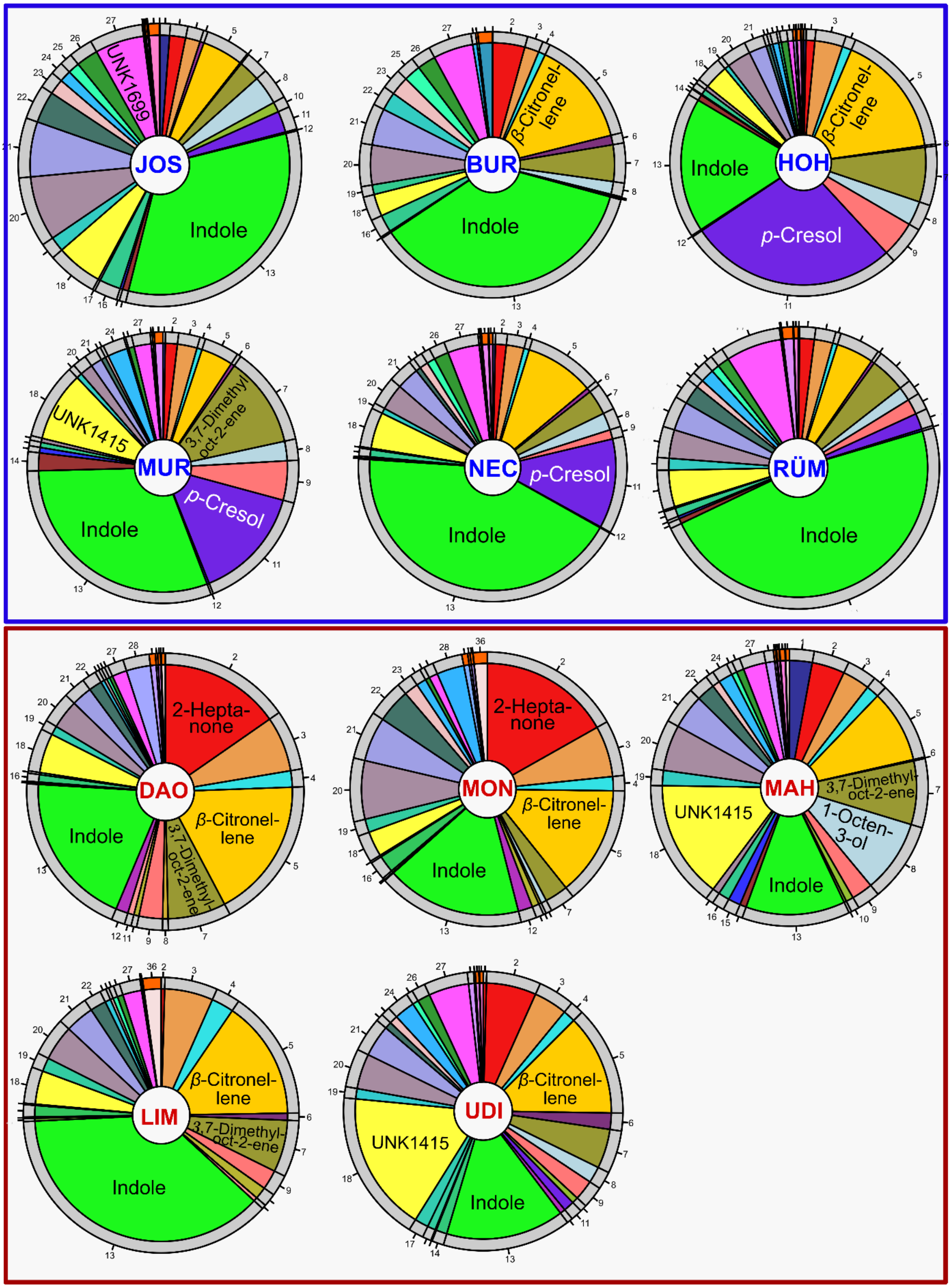
JOS
UNK1699
Indole
BUR
β-Citronel-lene
Indole
HOH
β-Citronel-lene
Indole
p-Cresol
MUR
UNK1415
3,7-Dimethyl oct-2-ene
p-Cresol
Indole
NEC
p-Cresol
Indole
RÜM
Indole
DAO
2-Hepta-none
β-Citronel-lene
3,7-Dimethyl-oct-2-ene
Indole
MON
2-Hepta-none
β-Citronel-lene
Indole
MAH
3,7-Dimethyl-oct-2-ene
1-Octen-3-ol
UNK1415
Indole
LIM
β-Citronel-lene
3,7-Dimethyl-oct-2-ene
Indole
UDI
β-Citronel-lene
UNK1415
Indole

**Fig. S2** Non-metric multidimensional scaling (nDMS) based on a Bray-Curtis dissimilarity matrix of relative (a, b) and absolute (c, d) scent patterns in *Arum maculatum*, using only compounds that correlated with relative fruit set (nine compounds; in a and c) or those that did not (93 compounds; in b and d; see also Material and Methods, Methods S3). Samples were collected in the two most extensively sampled populations north (JOS, blue) and south (DAO, red) of the Alps. See Table S1 for further identification of population codes.

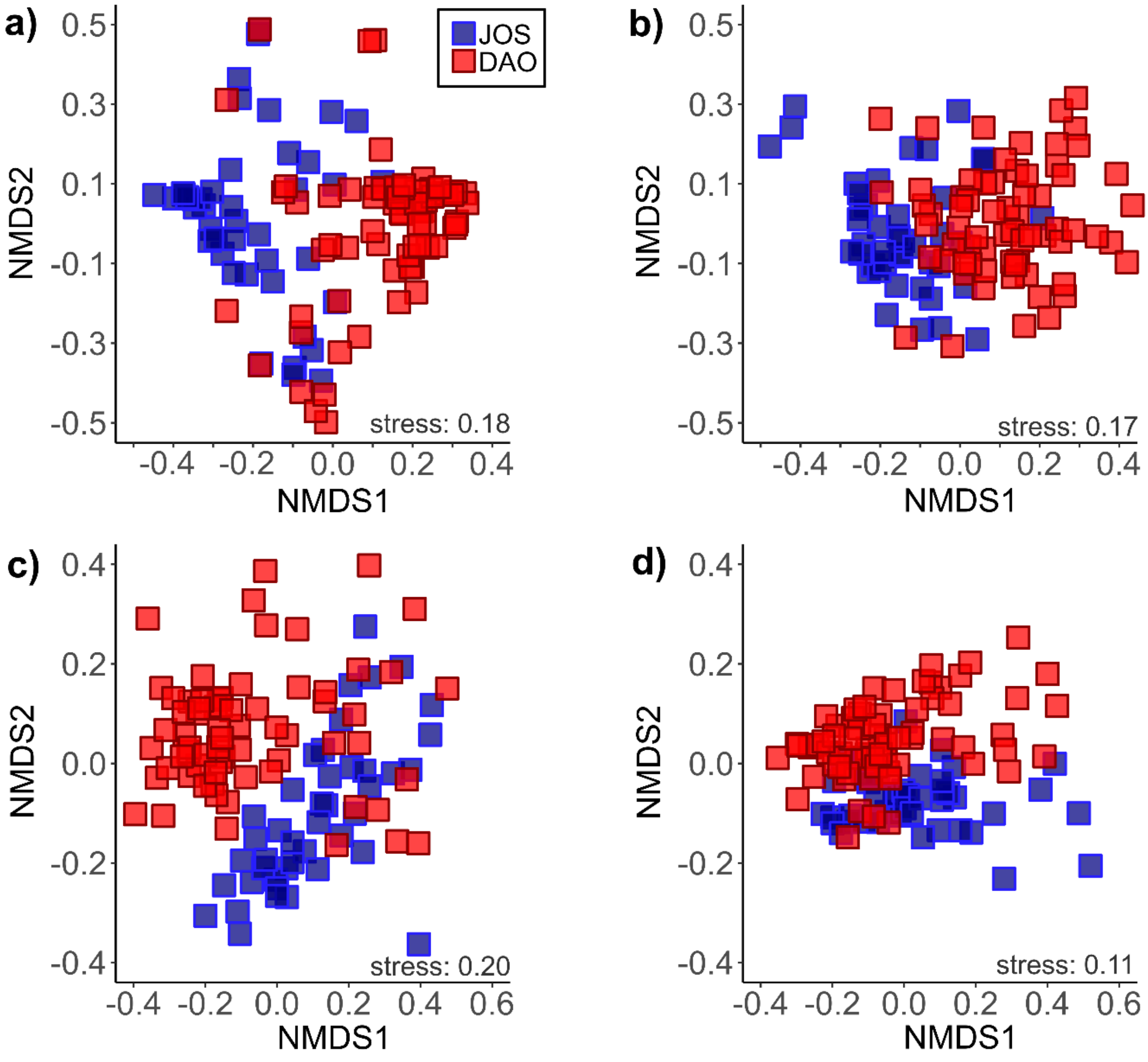

**Table S1** Locality information on the *Arum maculatum* populations sampled north (N) and south (S) of the Alps, including information on their altitudes.

| Code | Location | Country* | Altitude [m a.s.l.] | Latitude (N) | Longitude (E) |
|---|---|---|---|---|---|
| **N** | **North of the Alps** | | | | |
| JOS | Salzburg Josefiau | AT | 430 | 47° 46.98’ | 13° 04.50’ |
| BUR | Burg Hohenstein | D | 340 | 50° 11.64’ | 8° 03.42’ |
| HOH | Hohendilching | D | 630 | 47° 54.50’ | 11° 46.87’ |
| MUR | Murnau/Staffelsee | D | 660 | 47° 40.20’ | 11° 10.80’ |
| NEC | Horb/Neckar | D | 530 | 48° 25.20’ | 8° 39.00’ |
| RÜM | Rümikon/Aargau | CH | 350 | 47° 31.00’ | 8° 21.00’ |
| **S** | **South of the Alps** | | | | |
| DAO | Daone | IT | 900 | 45° 57.60’ | 10° 34.80’ |
| LIM | Limone-Piemonte | IT | 1010 | 44° 12.60’ | 7° 34.20’ |
| MAH | Santa Maria Hoè | IT | 430 | 45° 44.90’ | 9° 22.00’ |
| MON | Montese | IT | 845 | 44° 16.29’ | 10° 56.42’ |
| UDI | Udine | IT | 200 | 46° 10.59’ | 13° 15.55’ |

*AT (Austria), D (Germany), CH (Switzerland), IT (Italy)

**Table S3** The 25 volatile compounds (VOCs) most important for regional distinction in relative and absolute scent bouquets of *Arum maculatum* from north vs. south of the Alps. For each VOC, the mean decrease accuracy, as obtained by the *importance* measurement in *randomForest*, is given. Subscript letters in parentheses indicate the region where the compound was more abundant (N, north; S, south). *NA* indicates that this volatile was not among the top 25 in absolute or relative scent.

| VOC | Mean decrease accuracy | |
|---|---|---|
| | Absolute | Relative |
| UNK966 | 43.94 $_{(S)}$ | 47.55 $_{(S)}$ |
| 2-Heptanol | 44.20 $_{(S)}$ | 45.14 $_{(S)}$ |
| 2-Heptanone | 44.95 $_{(S)}$ | 41.60 $_{(S)}$ |
| 2-Nonanone | 43.33 $_{(S)}$ | 36.20 $_{(S)}$ |
| *α*-Citronellene | 38.43 $_{(S)}$ | 40.72 $_{(S)}$ |
| Dihydromyrcenol | 40.90 $_{(S)}$ | 35.52 $_{(S)}$ |
| 3,7-Dimethyloctan-1-ol | 33.64 $_{(S)}$ | 38.25 $_{(S)}$ |
| 1-Pentadecanol | 29.08 $_{(N)}$ | 42.31 $_{(N)}$ |
| 3,7-Dimethyloct-1-ene | 38.91 $_{(S)}$ | 31.09 $_{(S)}$ |
| *β*-Citronellene | 33.57 $_{(S)}$ | 34.27 $_{(S)}$ |
| UNK1207_1 | 29.44 $_{(S)}$ | 29.14 $_{(S)}$ |
| UNK1503 | 29.12 $_{(N)}$ | 28.49 $_{(N)}$ |
| UNK1063 | 27.87 $_{(N)}$ | 25.12 $_{(N)}$ |
| 2-Nonanol | 27.64 $_{(S)}$ | 24.81 $_{(S)}$ |
| 2,3-Butanediol | 22.79 $_{(N)}$ | 28.33 $_{(N)}$ |
| 3-Hepten-2-one | 25.76 $_{(S)}$ | 23.88 $_{(S)}$ |
| Indole | 19.98 $_{(S)}$ | 28.14 $_{(N)}$ |
| 1-Hexanol | 25.92 $_{(S)}$ | 20.57 $_{(S)}$ |
| *α*-Cadinene | 24.45 $_{(N)}$ | 20.18 $_{(N)}$ |
| UNK1469_2 | 20.01 $_{(N)}$ | 21.35 $_{(N)}$ |
| 2,6-Dimethyloct-7-en-4-one | 27.20 $_{(S)}$ | *NA* |
| 6-Methylheptan-2-one | 25.79 $_{(S)}$ | *NA* |
| UNK1415 | 25.72 $_{(S)}$ | *NA* |
| UNK681 | 21.58 $_{(S)}$ | *NA* |
| UNK1409 | 21.09 $_{(S)}$ | *NA* |
| *δ*-Cadinene | *NA* | 31.56 $_{(S)}$ |
| *p*-Cresol | *NA* | 29.72 $_{(N)}$ |
| 3-Methylbutanoic acid | *NA* | 21.32 $_{(N)}$ |
| UNK1794 | *NA* | 21.29 $_{(N)}$ |
| UNK676 | *NA* | 20.45 $_{(N)}$ |

**Table S4** Volatile compounds (VOCs) of *Arum maculatum* most correlating with the axes of the CAP ordination, based on absolute data (for more information see Materials and Methods).

| VOC | CAP1 [*r*] | CAP2 [*r*] |
|---|---|---|
| 1-Hexanol | 0.54*** | 0.07 *ns* |
| 2-Heptanol | 0.51*** | 0.18 *ns* |
| 2-Heptanone | 0.53*** | 0.21 * |
| 2-Nonanol | 0.51*** | 0.08 *ns* |
| 3,7-Dimethyloct-1-ene | 0.50*** | 0.11 *ns* |
| $\alpha$-Citronellene | 0.56*** | 0.05 *ns* |
| $\alpha$-Copaene | 0.54*** | -0.14 *ns* |
| $\alpha$-Humulene | 0.51*** | -0.10 *ns* |
| $\beta$-Caryophyllene | 0.56*** | -0.10 *ns* |
| $\beta$-Citronellene | 0.65*** | 0.09 *ns* |
| Indole | 0.65*** | -0.22 * |
| Isocaryophyllene | 0.54*** | -0.10 *ns* |
| UNK1164 | 0.53*** | -0.05 *ns* |
| UNK1378 | 0.58*** | -0.11 *ns* |
| UNK1391 | 0.58*** | -0.08 *ns* |
| UNK1403 | 0.56*** | -0.10 *ns* |
| UNK1424 | 0.58*** | -0.09 *ns* |
| UNK1438_1 | 0.54*** | -0.12 *ns* |
| UNK1457 | 0.61*** | -0.14 *ns* |
| UNK1466_1 | 0.53*** | -0.10 *ns* |
| UNK1492_1 | 0.53*** | -0.12 *ns* |

Significant values are marked with: $^{*}$, $P < 0.05$; $^{**}$, $P < 0.01$; $^{***}$, $P < 0.001$; *ns*, not significant.

**Table S5** Phenotypic selection (*β*- and *γ*- gradients) on volatile compounds (VOCs) of the two most extensively sampled *Arum maculatum* populations from north (JOS, model $N_{\beta}$) and south (DAO, model $S_{\gamma}$) of the Alps. Mann-Whitney-U-Tests (M-U-Test) indicate whether compounds differed in their absolute amounts between JOS and DAO, and the two regions (N, S). Compounds with significant gradients are in bold.

| VOC | Model | $\beta \pm$ SE | *P* | M-U-Test | |
|---|---|---|---|---|---|
| | | | | JOS vs. DAO | N vs. S |
| ***α*-Terpinene** | $N_{\beta}$ | 0.39±0.17 | 0.03 * | ** | *ns* |
| *β*-Lutidine | $N_{\beta}$ | -0.04±0.17 | 0.82 *ns* | *ns* | *ns* |
| Bicycloelemene | $N_{\beta}$ | 0.02±0.18 | 0.93 *ns* | ** | *ns* |
| Isobicyclogermacrene | $N_{\beta}$ | -0.30±0.11 | 0.11 *ns* | *ns* | *ns* |
| 1-Octanol | $N_{\beta}$ | 0.26±0.15 | 0.11 *ns* | *ns* | *ns* |
| 2,6-Dimethyl-7-octen-4-one | $N_{\beta}$ | 0.04±0.15 | 0.81 *ns* | * | *** |
| 2-Decanone | $N_{\beta}$ | -0.21±0.17 | 0.23 *ns* | *ns* | *ns* |
| **2-Heptanol** | $N_{\beta}$ | 0.34±0.14 | 0.02 * | *** | *** |
| **2-Nonanol** | $N_{\beta}$ | 0.37±0.18 | 0.04 * | *** | *** |
| **UNK681** | $N_{\beta}$ | 0.51±0.10 | < 0.001*** | *ns* | *** |
| UNK883 | $N_{\beta}$ | 0.29±0.15 | 0.07 *ns* | *** | ** |
| UNK960 | $N_{\beta}$ | -0.32±0.15 | 0.02 * | *ns* | *ns* |
| UNK1279 : 3-Octanol ‡ | $N_{\beta}$ | 0.23±0.12 | 0.07 *ns* | * : *** | ** : *ns* |
| UNK1409_1 | $N_{\beta}$ | -0.22±0.15 | 0.17 *ns* | *ns* | *** |
| UNK1440 | $N_{\beta}$ | 0.06±0.13 | 0.66 *ns* | *ns* | *** |
| **UNK1496 : UNK1503‡** | $N_{\beta}$ | 0.25±0.11 | 0.03 * | *ns : ** | *ns : ** |
| UNK1757_1 | $N_{\beta}$ | -0.14±0.14 | 0.31 *ns* | *ns* | *ns* |
| VOC | | $\gamma \pm$ SE | *P* | | |
| 2,6-Dimethylocta-2,6-diene | $S_{\gamma}$ | -0.04±0.25 | 0.88 *ns* | *** | *** |
| **4-Terpinenol** | $S_{\gamma}$ | -0.89±0.32 | 0.007** | * | *ns* |
| **Sabinene** | $S_{\gamma}$ | 8.96±1.77 | < 0.001*** | * | *** |

*, $P < 0.05$; **, $P < 0.01$; ***, $P < 0.001$; *ns*, not significant.

‡These VOCs were included in the $N_{\beta}$ model as a paired interaction term to avoid multicollinearity (for details see Materials and Methods).

**Methods S1** Detailed description of the scent analyses [thermal desorption-gas chromatography/mass spectrometry (TD-GC/MS), quantification and building of a scent library for semi-automatic analysis].

We used a thermal desorption system (model TD-20, Shimadzu, Tokyo, Japan) coupled with a GC/MS (QP2010 Ultra El, Shimadzu), and equipped with a ZB-5 fused silica column (5% phenyl polysiloxane; 60 m long, inner diameter 0.25 mm, film thickness 0.25 µm, Phenomenex, Torrance, CA, USA). The column flow (carrier gas: helium) was set to 1.5 ml $min^{-1}$, at a split ratio between 1 : 1 and 1 : 5, except for a single sample that was analysed at a split ratio of 1 : 20. The split ratio was set depending on the perceived strength of the scent emitted by the plants used for scent collections. Starting at 40 °C, the GC oven temperature was increased by 6 °C per min to 250 °C and was held for 1 min. The MS interface worked at 250 °C and mass spectra were taken at 70 eV from *m/z* 30 to 350 (i.e., mass-to-charge ratio, in EI mode). The GC/MS data were handled using the GCMSolution package version 4.41 (Shimadzu Corporation, Kyoto, Japan). We established an own library of mass-spectral and Kováts' retention indices (KRIs[1]; based on *n*-alkanes series ($C_7$–$C_{20}$)) for semi-automatic analysis. Therefore, we manually analysed five indidvidual samples per population and included all detected inflorescence-specific compounds in the new library. We then manually scanned all other individual scent samples for additional, not yet included, floral compounds to complete the library with our target compounds. Afterwards, we automatically searched for and integrated the target compounds available in the chromatograms and, whenever necessary, completed this by manually integrating peaks not automatically detected (e.g., due to coelution with other compounds). To estimate the total absolute emission of scent trapped while considering that samples were run at different split settings, known amounts of the *n*-alkane series (0.3–1.0 µl) were injected at these split settings, and the resulting mean peak areas of the central alkanes $C_{14}$–$C_{16}$ were used for quantification.

**Methods S2** Synthesis of reference samples: 2,6-dimethylocta-2,6-diene, 3,7-dimethyloct-2-ene and 2,6-dimethylocta-1,7-diene ($\alpha$-citronellene).

2,6-Dimethylocta-2,6-diene: A suspension of 11.18 g (30.11 mmol) ethyltriphenylphosphonium bromide in 120 ml dry THF under argon was cooled to -40°C and *n*-butyllithium solution (1.6 M in hexanes) was added dropwise, until the colour of the mixture remained slightly yellow (0.5 ml). Subsequently, additional 18.82 ml BuLi-solution (30.11 mmol) was carefully added, turning the colour of the mixture to dark orange. After stirring for another hour at -35°C, a solution of 3.78 g (30 mmol) 6-methyl-5-hepten-2-one (sulcatone) in THF (10 ml) was added and the cooling bath removed. After reaching room temperature, the mixture was poured into ice water (200 ml) and extracted with *n*-hexane (3 × 150 ml). The organic extracts were combined, washed with water and brine, dried over $MgSO_4$ and concentrated under reduced pressure. Subsequent purification by flash chromatography (40 g $SiO_2$, *n*-pentane) gave 3.18 g 2,6-dimethyl-2,6-octadiene (78 % yield) as a colorless oil.

3,7-Dimethyloct-2-ene: Using the same standard procedure (Wittig procedure) as described above, 7.99 g (21.5 mmol) ethyltriphenylphosphonium bromide, 13.44 ml (21.5 mmol) *n*-butyllithium solution and 2.52 g (19.68 mmol) 6-methyl-2-heptanone yielded 1.98 g (14.14 mmol, 72%) of the desired 3,7-dimethyl-2-octene.

2,6-Dimethylocta-1,7-diene ($\alpha$-citronellene): According to[2] 10 mg *p*-toluenesulfonic acid was added to a sample of (6S)-3,7-dimethyl-1,6-octadiene (2.0 g, 14.5 mmol) and the mixture was stirred for 12 h at 110°C under argon. Subsequent distillation using a 10 cm Vigreux column (45 mbar, ~72°C) yielded 1.3 g (9.4 mmol) of a 3 : 1 mixture of' the educt and the desired 2,6-dimethyl-1,7-octadiene. This was used as a reference sample without further purification.

**Methods S3** Simulation study to quantify impact of non-detects (zero-inflation) on selection estimates and reducing high dimensionality of scent data by determining scent compounds that correlate with fruit set via elastic net (linear) and *Boruta* (nonlinear) analyses.

Simulation: To quantify the impact of non-detects on estimates, we performed a simulation study based on our scent data for each of the most extensively sampled population (JOS, DAO) separately. For these simulations, we first considered per dataset a subset of variables (compounds) with at least five non-zero values, to ensure variability in the sampled data and so that the number of zeros could be controlled for the simulation. This resulted in 209 and 213 variables for the most extensively sampled northern (JOS) and southern (DAO), respectively. We then sampled from these variables with replacement, constructed an elastic net model[1] based on these predictors with $\alpha$ ($L_1/L_2$ penalty ratio) set at 0 ($L_1$, lasso), 0.5 ($L_1/L_2$), or 1 ($L_2$, ridge), and $\lambda$ (penalty strength parameter), cross-validated for each $\alpha$ (see also below), and set the smallest x% of values to zero, where x ranged from 30% to 60%. With the assumption that either 50% or 10% of the variables are important predictors for the model, we then assessed how the sensitivity and specificity of the full vs. zero-inflated models changed. All simulations were repeated 10,000 times. To keep the influence of zero-inflation at a maximum of 5%, i.e., the difference of sensitivity and specificity between the full and zero-inflated model, we decided to set the threshold to a maximum of 50% non-detects in both populations, even though the obtained differences varied between the two populations. This resulted in 93 and 81 scent variables for JOS and DAO, respectively. The R code for this simulation [will be] available at the Dryad digital repository.

Elastic Net: An elastic net regression shrinks coefficients that are not important for the model towards zero ($L_1$ penalisation) by adding a penalty to the regression (regularisation). Additionally, by $L_2$ penalisation, it selects groups of correlated variables are selected instead of only one random variable from a multicollinear group of variables (such as a LASSO regression does)[3]. By combining $L_1$ with $L_2$ penalisation, elastic net regression is well suited for datasets that have a larger number of predictor variables (in our case scent compounds) than number of observations, and copes well with multicollinearity[3,4]. The latter is of particular advantage for our study as volatiles sharing the same biosynthetic pathway can be highly multicollinear. Zero-inflation can still influence these analyses, yet this was solved by the simulation (see above). For the elastic net, we first cross-validated for $\alpha$ ($L_1/L_2$ penalty ratio, range: 0−1, 0.1 increment) and $\lambda$ (penalty strength parameter; range: 100−0.001, -0.1 increment), for the reduced datasets of JOS and DAO (see above) separately. The final elastic net (family = "gaussian") for the northern JOS population was run with $\alpha_{(\mathrm{JOS})} = 0.9$ and for the southern population DAO with $\alpha_{(\mathrm{DAO})} = 0.2$, and regression coefficients $\hat{\beta}$ were extracted at $\lambda_{min(\mathrm{JOS})} = 0.05$ and $\lambda_{min(\mathrm{DAO})} = 1.99$. For all elastic net analyses, we used the R packages *glmnet* v.4.1[10].

Boruta: Pollinators might respond to different amounts of scent compounds also in a nonlinear manner[5–7], potentially resulting in a nonlinear relationship of fitness and scent traits. To account for this, we ran a *Boruta* algorithm (R package *Boruta* v.7.0.0[8]; *ntree* = 9,999 bootstrap samples with *mtry* = 9, Bonferroni-adjusted for multiple comparisons) to also identify nonlinear relationships between individual volatiles, total absolute scent emission, and relative fruit set. *Boruta* is a feature selection method that has shown high stability in variable selection of high-dimensional data sets[9] and classifies whether a feature (in our case scent compounds) is important or not[8].

**Notes S1** Mass spectra of unknown volatiles with significant phenotypic selection gradients.

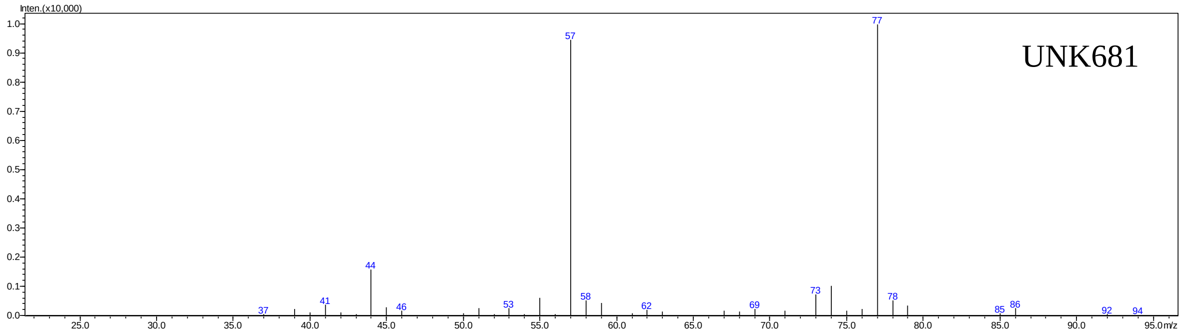

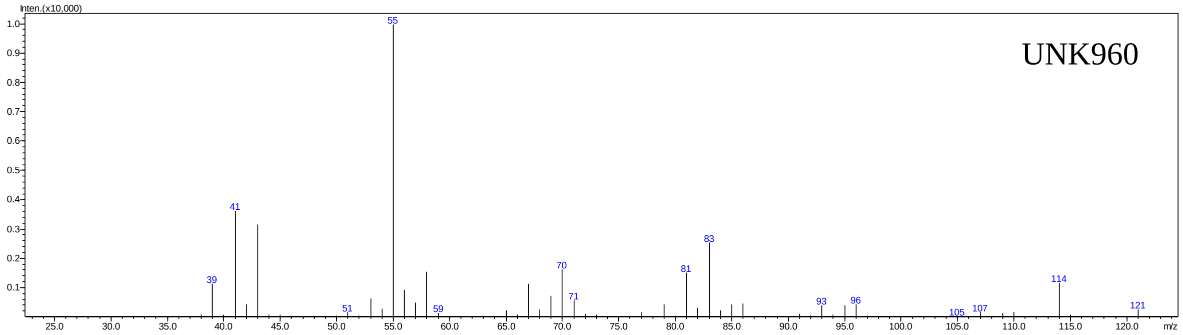

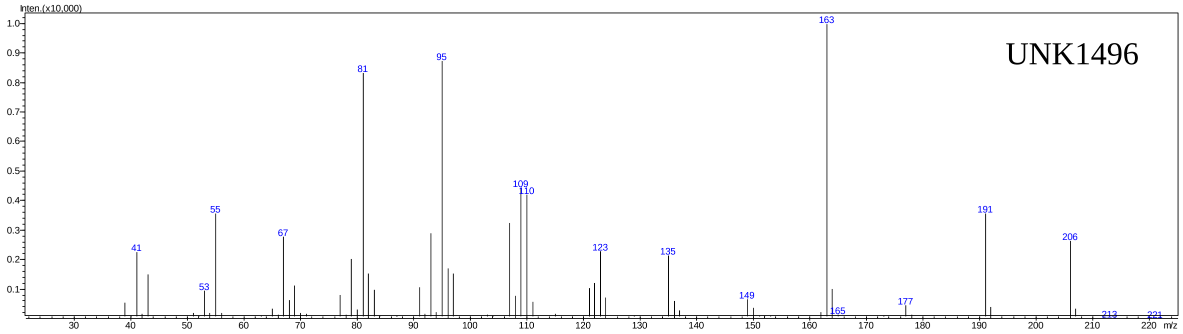

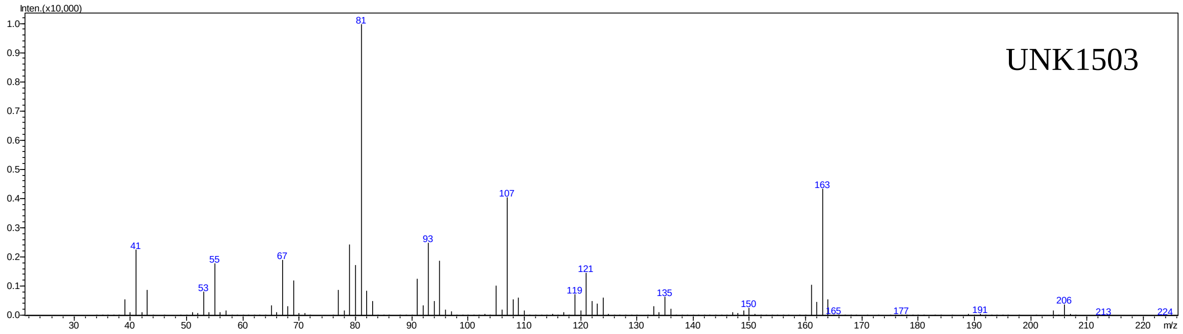
