## Supporting Information Table S2 for "Floral scents of a deceptive plant are hyperdiverse and under population-specific phenotypic selection"

**Table S2** Median amounts of total absolute and relative (contribution of single compounds to total scent) inflorescence scent of *Arum maculatum* , surveyed in six and five populations north and south of the Alps, respectively. The total number of volatiles is also given.

[a]KRI, Kováts' retention index compounds in **bold** are >1% in at least one population tr = trace relative amount <0.05% – = compound did not occur in any individual

* = identification of compound was verified by authentic standards. Underlined compounds indicate compounds under selection (see Fig. 5). See Table S1 for identification of population codes

| KRI[a] | | North | JOS | BUR | HOH | MUR | NEC | RÜM | South | DAO | LIM | MAH | MON | UDI |
|---|---|---|---|---|---|---|---|---|---|---|---|---|---|---|
| | Median total absolute emission of scent (ng inflorescence$^{-1}$ h$^{-1}$) | 67.4 | 167.2 | 13 | 565.8 | 80.7 | 39.4 | 41.7 | 214.7 | 311.4 | 203.8 | 196.9 | 201.4 | 42.3 |
| | Total number of volatiles | 283 | 269 | 166 | 197 | 204 | 181 | 208 | 265 | 254 | 195 | 146 | 88 | 171 |
| **Aliphatic components** | | | | | | | | | | | | | | |
| 707 | Acetoin* | tr | tr | – | – | tr | – | tr | tr | tr | – | tr | – | tr |
| 753 | 3-Methylpentan-2-one* | tr | 0.1 | tr | – | tr | tr | tr | tr | tr | – | tr | tr | tr |
| 776 | Butanoic acid* | tr | tr | – | 0.9 | tr | 0.6 | tr | tr | tr | 0.3 | tr | tr | – |
| 787 | 2,3-Butanediol* | tr | tr | tr | – | 0.2 | tr | tr | tr | tr | – | tr | tr | – |
| 870 | 1-Hexanol* | 0.2 | 0.2 | 0.1 | 0.3 | 0.2 | 0.2 | 0.1 | 0.4 | 0.4 | 0.5 | 0.3 | 0.3 | 0.4 |
| 885 | 2,3-Heptanedione | tr | tr | – | – | – | – | – | tr | tr | – | – | – | – |
| 893 | **2-Heptanone*** | 1.4 | 1.4 | 2.4 | 0.8 | 1.3 | 1.1 | 1.4 | 6.9 | 9.3 | 0.3 | 2.9 | 11.9 | 4 |
| 902 | **2-Heptanol*** | 0.1 | 0.1 | 0.3 | 0.3 | 0.2 | 0.2 | 0.1 | 1.2 | 1.9 | tr | 0.8 | 2.5 | 0.5 |
| 926 | Methyl hexanoate* | tr | tr | tr | 0.2 | tr | tr | tr | tr | tr | – | tr | – | tr |
| 938 | 3-Hepten-2-one | tr | tr | tr | tr | – | tr | – | tr | tr | tr | – | tr | tr |
| 939 | Methyl 4-methylpent-4-eneoate | tr | tr | – | tr | tr | tr | – | tr | – | – | tr | – | – |
| 947 | 4-Methylhexan-1-ol | tr | tr | tr | tr | tr | tr | tr | tr | tr | tr | tr | tr | tr |
| 953 | Isobutyl butyrate | tr | tr | – | 0.1 | – | tr | – | tr | – | – | tr | – | – |
| 982 | **1-Octen-3-ol*** | 1.9 | 2.4 | tr | 2.3 | 2 | 1.8 | 1.3 | 0.3 | 0.4 | tr | 6.4 | tr | 1.3 |
| 989 | 3-Octanone* | tr | tr | – | – | 0.6 | tr | tr | 0.3 | 0.4 | – | 0.8 | tr | 0.2 |
| 998 | 3-Octanol* | tr | tr | tr | – | 0.1 | tr | tr | tr | tr | – | 0.3 | – | tr |
| 1069 | 1-Octanol* | tr | tr | tr | – | tr | tr | tr | tr | tr | – | tr | – | tr |
| 1071 | (E)-2-Octen-1-ol* | tr | tr | tr | tr | tr | tr | – | tr | tr | – | 0.2 | tr | – |
| 1096 | **2-Nonanone*** | 0.2 | 0.1 | 0.1 | 0.1 | 0.2 | 0.1 | 0.2 | 0.7 | 0.9 | tr | 0.2 | 1.3 | 0.4 |
| 1099 | 2-Nonanol* | 0.1 | 0.1 | tr | tr | 0.2 | 0.1 | tr | 0.2 | 0.4 | tr | 0.2 | 0.5 | 0.1 |
| 1123 | Methyl octanoate* | tr | tr | tr | – | tr | tr | – | – | – | – | – | – | – |
| 1160 | 2-Decanone | 0.1 | tr | 0.1 | 0.1 | 0.1 | 0.1 | 0.1 | tr | tr | tr | tr | 0.1 | 0.1 |
| 1294 | 2-Undecanone | tr | tr | tr | tr | tr | tr | tr | tr | tr | tr | tr | tr | tr |
| 1303 | 2-Undecanol* | tr | tr | – | – | tr | tr | tr | tr | tr | – | tr | tr | – |
| 1538 | 1-Tridecanol | tr | tr | – | – | tr | tr | tr | tr | tr | – | – | – | – |
| 1745 | 1-Pentadecanol | 0.2 | 0.1 | 0.3 | 0.3 | 0.2 | 0.3 | 0.3 | tr | tr | 0.1 | tr | 0.1 | tr |
| **Aromatic components** | | | | | | | | | | | | | | |
| 1026 | 4-Methylanisol* | tr | tr | – | – | – | – | – | tr | – | tr | tr | tr | – |
| 1076 | ***p*-Cresol*** | 4.2 | 1.8 | 0.1 | 19.4 | 11.9 | 9.2 | 1.5 | 0.5 | 0.5 | 0.3 | 0.7 | 0.6 | 0.9 |
| 1199 | *p* -Creosol | tr | tr | – | tr | 0.1 | tr | tr | tr | tr | tr | tr | – | tr |
| 1205 | Methyl salicylate* | tr | tr | tr | 0.5 | 0.2 | 0.1 | 0.1 | tr | tr | tr | 0.1 | tr | tr |
| 1367 | *p* -Cresyl butyrate | tr | – | – | tr | – | tr | – | – | – | – | – | – | – |
| **C5-branched chain components** | | | | | | | | | | | | | | |
| 780 | Methyl 2-methylbutyrate* | tr | tr | – | tr | tr | – | – | tr | tr | – | tr | tr | – |
| 831 | 3-Methylbutanoic acid* | tr | tr | tr | tr | tr | 0.1 | 0.1 | tr | tr | – | tr | – | tr |
| 842 | 2-Methylbutanoic acid* | tr | tr | tr | 0.1 | tr | 0.1 | tr | tr | tr | – | tr | – | tr |
| 1105 | Isoamyl isovalerate | tr | tr | tr | – | – | – | tr | tr | tr | tr | – | tr | tr |
| **Nitrogen-bearing components** | | | | | | | | | | | | | | |

| | | | | | | | | | | | | | | |
|---|---|---|---|---|---|---|---|---|---|---|---|---|---|---|
| 965 | **β-Lutidine** | 0.2 | 0.1 | 0.7 | 0.2 | 0.4 | 0.4 | 0.3 | 0.1 | tr | 0.6 | 0.1 | tr | 1.3 |
| 1232 | 2-Aminobenzaldehyde* | tr | tr | – | – | – | tr | tr | tr | tr | tr | – | – | – |
| 1310 | **Indole*** | 24.2 | 22.3 | 20.8 | 12.6 | 24.6 | 33.4 | 35.6 | 11.9 | 11.9 | 24.8 | 8.8 | 12.3 | 9.5 |
| 1314 | 2-Aminoacetophenone* | tr | tr | – | – | – | tr | – | tr | tr | tr | tr | – | – |
| 1360 | Methyl anthranilate* | tr | tr | – | – | – | – | – | tr | tr | tr | – | tr | tr |
| 1406 | Skatole* | 0.1 | 0.1 | tr | 0.1 | tr | tr | 0.3 | 0.1 | tr | 0.1 | 0.1 | tr | tr |
| 1428 | 2-(Dimethylamino)benzaldehyde | tr | tr | – | – | – | tr | tr | – | – | – | – | – | – |
| **Irregular terpene** | | | | | | | | | | | | | | |
| 959 | 6-Methylheptan-2-one | tr | tr | 0.1 | tr | tr | 0.1 | 0.1 | 0.1 | 0.1 | tr | tr | 0.3 | tr |
| 992 | 6-Methylhept-5-ene-2-ol | tr | tr | tr | 0.3 | tr | 0.4 | tr | tr | tr | tr | tr | tr | tr |
| 1853 | Hexahydrofarnesylacetone* | tr | tr | 0.3 | 0.1 | tr | 0.1 | tr | tr | tr | 0.2 | – | tr | 0.1 |
| **Monoterpenoids** | | | | | | | | | | | | | | |
| 914 | **3,7-Dimethyloct-1-ene*** | 1.5 | 1.2 | 1.1 | 2.6 | 2.1 | 1.8 | 1.7 | 4 | 4.3 | 4.1 | 2.4 | 4.6 | 2.7 |
| 935 | **α-Citronellene***[§] | 0.4 | 0.3 | 0.5 | 0.9 | 0.5 | 0.5 | 0.4 | 1.3 | 1.3 | 1.9 | 1.1 | 1.3 | 1 |
| 949 | **β- Citronellene***[§] | 4.2 | 3.5 | 8 | 11.6 | 3.4 | 7.1 | 3.5 | 9.7 | 10.8 | 10.1 | 6.5 | 9.9 | 8.2 |
| 996 | β-Myrcene* | tr | tr | tr | 0.4 | tr | tr | tr | tr | tr | tr | tr | tr | tr |
| 972 | **3,7-Dimethyloct-2-ene*** | 3.1 | 1.8 | 2.8 | 5.1 | 9.6 | 2.6 | 3.3 | 4.3 | 4.5 | 4.4 | 5.6 | 2.1 | 3.2 |
| 982 | **<u>Sabinene</u>*** | 0.2 | tr | 1 | 0.2 | tr | 0.4 | 0.4 | 0.4 | 0.3 | 1.4 | 0.3 | 1.2 | tr |
| 1005 | **2,6-Dimethylocta-2,6-diene*** | tr | tr | tr | tr | tr | tr | tr | 0.4 | 0.4 | 1 | tr | 0.2 | 0.5 |
| 1025 | <u>α-Terpinene*</u> | tr | tr | tr | tr | tr | tr | tr | tr | tr | tr | 0.1 | – | tr |
| 1039 | β-Phellandrene* | tr | tr | tr | tr | tr | tr | tr | tr | tr | tr | 0.1 | tr | tr |
| 1050 | 2,6-Dimethylocta-3,7-dien-2-ol | tr | tr | tr | – | 0.1 | tr | 0.1 | tr | tr | tr | tr | 0.1 | – |
| 1051 | (E)-β-Ocimene* | tr | tr | tr | 0.1 | tr | 0.1 | tr | tr | tr | 0.2 | tr | tr | tr |
| 1058 | 2,6-Dimethyloct-7-en-4-one | tr | tr | tr | – | 0.1 | 0.1 | tr | tr | tr | 0.2 | 0.1 | tr | tr |
| 1068 | γ-Terpinene* | tr | 0.1 | tr | 0.1 | tr | tr | tr | tr | tr | tr | 0.2 | tr | tr |
| 1074 | (Z)-Sabinenehydrate | tr | – | tr | – | – | – | tr | tr | tr | tr | tr | tr | tr |
| 1076 | **Dihydromyrcenol** | tr | tr | tr | tr | tr | tr | tr | 0.4 | 0.4 | 1 | tr | 0.2 | 0.5 |
| 1096 | α-Terpinolene* | tr | tr | tr | 0.1 | – | tr | – | tr | tr | 0.2 | 0.1 | tr | – |
| 1100 | Linalool* | tr | tr | tr | 0.3 | 0.1 | 0.1 | 0.2 | tr | tr | 0.1 | tr | tr | 0.1 |
| 1122 | Myrcenol | tr | – | – | tr | – | – | – | tr | tr | tr | – | – | – |
| 1135 | *allo*-Ocimene* | tr | tr | tr | 0.2 | – | 0.1 | tr | tr | tr | tr | tr | tr | – |
| 1149 | *neoallo*-Ocimene* | tr | tr | tr | 0.1 | 0.1 | tr | tr | tr | tr | 0.1 | tr | tr | tr |
| 1157 | Citronellal* | tr | tr | – | tr | – | tr | tr | tr | tr | tr | tr | tr | tr |
| 1179 | 2,7-dimethylocta-4,6-dien-2-ol | tr | tr | tr | 0.2 | 0.2 | tr | 0.1 | tr | tr | 0.2 | tr | tr | tr |
| 1191 | <u>4-Terpinenol</u>* | tr | tr | tr | tr | tr | 0.1 | 0.1 | tr | tr | 0.1 | tr | tr | tr |
| 1195 | *p*-Cymen-8-ol | tr | tr | tr | tr | tr | tr | tr | tr | tr | 0.1 | tr | 0.1 | – |
| 1195 | 3,7-Dimethyloctan-1-ol | 0.1 | 0.1 | tr | 0.2 | 0.2 | 0.1 | 0.2 | 0.4 | 0.5 | 0.3 | 0.5 | 0.2 | 0.4 |
| 1201 | α-Terpineol* | tr | tr | tr | 0.1 | tr | tr | tr | tr | tr | tr | tr | tr | 0.1 |
| 1232 | Nerol* | tr | tr | tr | tr | tr | tr | tr | tr | tr | tr | tr | – | 0.2 |
| 1233 | β-Citronellol* | 0.1 | 0.1 | tr | 0.3 | 0.2 | 0.1 | 0.3 | 0.1 | 0.1 | 0.3 | 0.1 | tr | 0.2 |
| 1255 | Carvone* | tr | tr | – | – | – | – | – | tr | tr | tr | – | – | – |
| **Sesquiterpenoids** | | | | | | | | | | | | | | |
| 1346 | Silphiperfol-5-ene | – | – | – | – | – | – | – | tr | tr | – | – | – | – |
| 1357 | **Bicycloelemene** | 0.4 | 0.5 | 0.1 | 0.5 | 1.9 | 0.1 | 0.5 | 0.2 | 0.1 | 0.2 | 0.6 | 0.1 | 0.9 |
| 1366 | α-Cubebene | tr | tr | tr | tr | – | tr | tr | tr | tr | tr | – | – | – |
| 1399 | **α-Copaene*** | 1 | 1.8 | 1.3 | 0.5 | 0.5 | 0.7 | 0.8 | 0.8 | 0.6 | 1 | 1 | 1.6 | 0.6 |
| 1434 | **Isocaryophyllene** | 0.9 | 1.3 | 0.7 | 0.4 | 0.6 | 0.5 | 1.1 | 0.9 | 0.7 | 1.2 | 1.3 | 1.2 | 0.8 |
| 1450 | **β-Caryophyllene*** | 3 | 5.5 | 3 | 2.3 | 1.4 | 2.7 | 2.7 | 2.9 | 2.2 | 3 | 4 | 5.3 | 2.8 |

| | | | | | | | | | | | | | | |
|---|---|---|---|---|---|---|---|---|---|---|---|---|---|---|
| 1484 | ***α*-Humulene*** | 2.8 | 4.7 | 2.8 | 1.7 | 1.2 | 2.3 | 2.7 | 2.3 | 1.6 | 2.5 | 3 | 4 | 2.6 |
| 1501 | **Germacrene D*** | 0.9 | 1.3 | 1.4 | 0.3 | 0.3 | 0.9 | 0.7 | 0.5 | 0.3 | 0.5 | 0.7 | 1.3 | 0.7 |
| 1505 | Isobicyclogermacrene | tr | tr | tr | 0.1 | 0.1 | tr | 0.1 | tr | tr | tr | 0.1 | tr | 0.1 |
| 1513 | (E,E)-*α*-Farnesene* | tr | tr | – | tr | tr | – | tr | tr | tr | tr | tr | – | tr |
| 1520 | **Bicyclogermacrene** | 0.9 | 1 | tr | 0.6 | 2.1 | tr | 1.3 | 0.4 | 0.2 | 0.2 | 1.3 | tr | 1.7 |
| 1528 | (*E*)-Calamene | tr | tr | – | – | tr | tr | – | tr | – | – | – | – | tr |
| 1547 | ***δ*-Cadinene** | 1.2 | 1.9 | 1.4 | 0.6 | 0.6 | 1.5 | 1.1 | 0.4 | 0.3 | 0.5 | 0.7 | 0.4 | 1 |
| 1550 | (*Z*)-Calamene | 0.1 | 0.1 | tr | tr | tr | tr | 0.1 | tr | tr | 0.1 | 0.1 | 0.1 | tr |
| 1564 | *α*-Cadinene | tr | 0.1 | tr | tr | tr | tr | 0.1 | tr | tr | tr | tr | tr | tr |
| 1572 | *α*-Calacorene | 0.2 | 0.2 | 0.2 | 0.1 | 0.1 | 0.2 | 0.2 | 0.1 | 0.1 | 0.1 | 0.1 | 0.2 | 0.1 |
| 1592 | *β*-Calacorene | 0.1 | 0.1 | 0.1 | tr | 0.1 | 0.1 | 0.1 | 0.1 | tr | 0.1 | 0.1 | 0.2 | 0.1 |
| 1707 | Cadalene | tr | 0.1 | tr | 0.1 | tr | 0.1 | tr | tr | tr | tr | 0.1 | 0.1 | tr |
| **Unknown compounds** | | | | | | | | | | | | | | |
| 676 | UNK 676 | tr | tr | 0.1 | tr | tr | tr | tr | tr | tr | tr | tr | tr | tr |
| 681 | <u>UNK 681</u> | tr | 0.2 | tr | tr | tr | tr | tr | 0.2 | 0.2 | tr | 0.2 | 0.2 | 0.3 |
| 744 | UNK 744 | – | – | – | – | – | – | – | tr | tr | tr | – | – | – |
| 829 | **UNK 829** | 0.3 | 0.8 | tr | 0.2 | 0.2 | 0.3 | 0.1 | tr | tr | tr | 2 | tr | 0.3 |
| 849 | UNK 849 | tr | tr | tr | – | tr | tr | tr | tr | tr | tr | tr | tr | tr |
| 854 | UNK 854 | tr | tr | – | – | – | – | – | tr | tr | – | tr | – | – |
| 861 | UNK 861 | tr | tr | – | – | – | tr | tr | tr | – | – | – | – | tr |
| 863 | UNK 863 | tr | tr | – | – | – | – | – | tr | tr | tr | – | tr | tr |
| 883 | UNK 883 | tr | tr | tr | – | tr | tr | tr | tr | tr | – | tr | – | tr |
| 888 | UNK 888 | tr | tr | tr | tr | tr | tr | tr | tr | tr | tr | tr | tr | tr |
| 895 | UNK 895 | tr | tr | – | – | – | – | – | tr | tr | – | tr | – | tr |
| 922 | UNK 922 | tr | tr | tr | tr | tr | – | tr | tr | tr | – | tr | tr | tr |
| 925 | UNK 925 | tr | tr | – | tr | – | – | – | tr | – | tr | – | – | – |
| 938 | UNK 938 | tr | tr | – | – | tr | – | – | tr | tr | – | tr | – | tr |
| 960 | <u>UNK 960</u> | tr | tr | – | tr | – | – | – | tr | – | – | – | – | tr |
| 962 | UNK 962 | 0.1 | 0.1 | tr | 0.1 | 0.1 | 0.1 | 0.1 | 0.1 | 0.1 | 0.2 | 0.2 | 0.1 | tr |
| 966 | UNK 966 | tr | tr | tr | tr | tr | tr | tr | 0.2 | 0.2 | 0.5 | – | 0.3 | 0.2 |
| 978 | UNK 978 | tr | 0.1 | tr | 0.1 | tr | tr | tr | tr | tr | – | 0.1 | tr | tr |
| 991 | UNK 991 | tr | tr | – | tr | – | tr | – | tr | tr | tr | tr | – | – |
| 995 | UNK 995 | tr | tr | – | – | tr | – | tr | tr | tr | tr | – | – | – |
| 1005 | UNK 1005 | tr | tr | tr | tr | – | – | – | tr | tr | tr | tr | tr | tr |
| 1014 | UNK 1014 | tr | tr | tr | 0.1 | 0.1 | tr | tr | tr | tr | tr | tr | – | tr |
| 1021 | UNK 1021 | tr | tr | – | – | – | – | – | tr | tr | – | – | – | – |
| 1026 | UNK 1026 | tr | tr | – | tr | tr | – | tr | tr | tr | tr | tr | tr | – |
| 1030 | UNK 1030 | 0.2 | 0.2 | 0.1 | 0.7 | 0.8 | 0.1 | 0.3 | 0.1 | 0.1 | 0.3 | 0.2 | tr | 0.4 |
| 1032 | UNK 1032 | tr | tr | – | tr | – | tr | tr | tr | – | – | tr | – | tr |
| 1033 | UNK 1033 | tr | tr | tr | tr | tr | tr | tr | tr | tr | tr | – | – | tr |
| 1044 | UNK 1044 | tr | – | – | tr | – | – | – | tr | tr | – | – | – | tr |
| 1052 | UNK 1052 | tr | tr | tr | tr | tr | – | tr | tr | tr | tr | tr | – | tr |
| 1057 | UNK 1057 | tr | tr | – | – | – | – | – | tr | tr | – | – | tr | tr |
| 1058 | UNK 1058 | tr | tr | – | – | – | tr | – | – | – | – | – | – | – |
| 1059 | UNK 1059 | tr | tr | – | tr | – | – | – | tr | – | – | tr | – | tr |
| 1060 | UNK 1060 | tr | tr | – | tr | tr | tr | – | tr | – | – | tr | – | tr |
| 1063 | UNK 1063 | tr | tr | – | 0.2 | – | tr | tr | tr | tr | – | tr | – | – |
| 1064 | UNK 1064 | tr | tr | – | – | – | – | – | tr | tr | – | – | tr | – |

| | | | | | | | | | | | | | | |
|---|---|---|---|---|---|---|---|---|---|---|---|---|---|---|
| 1068 | UNK 1068_1 | 0.1 | tr | 0.1 | 0.3 | 0.2 | 0.1 | 0.2 | 0.1 | 0.1 | 0.3 | tr | 0.1 | 0.3 |
| 1068 | UNK 1068_2 | tr | tr | – | – | tr | – | tr | tr | tr | – | tr | – | tr |
| 1084 | UNK 1084 | tr | tr | – | tr | tr | tr | – | tr | tr | – | tr | tr | tr |
| 1085 | UNK 1085 | tr | tr | tr | tr | tr | tr | tr | tr | tr | tr | tr | – | tr |
| 1094 | UNK 1094 | tr | tr | tr | – | 0.2 | 0.1 | 0.2 | tr | tr | tr | tr | 0.1 | tr |
| 1105 | UNK 1105 | tr | tr | – | tr | – | tr | – | tr | tr | – | tr | tr | – |
| 1113 | UNK 1113 | tr | tr | tr | – | – | tr | tr | tr | – | – | – | – | tr |
| 1114 | UNK 1114 | tr | tr | tr | tr | tr | tr | tr | tr | tr | tr | tr | tr | – |
| 1124 | UNK 1124_1 | tr | tr | tr | tr | tr | tr | tr | tr | tr | tr | tr | tr | – |
| 1124 | UNK 1124_2 | tr | tr | tr | tr | tr | tr | – | tr | tr | tr | tr | tr | – |
| 1128 | UNK 1128 | tr | tr | tr | 0.1 | 0.1 | tr | tr | tr | tr | tr | tr | – | – |
| 1130 | UNK 1130 | tr | tr | tr | – | tr | – | – | tr | tr | tr | – | tr | – |
| 1134 | UNK 1134 | tr | – | tr | – | 0.1 | – | 0.1 | tr | tr | – | – | tr | – |
| 1135 | UNK 1135 | tr | tr | tr | 0.1 | 0.2 | tr | 0.1 | 0.1 | 0.1 | 0.4 | tr | tr | 0.1 |
| 1139 | UNK 1139 | tr | tr | tr | tr | tr | tr | tr | tr | tr | tr | tr | tr | tr |
| 1146 | UNK 1146 | tr | tr | tr | tr | 0.1 | tr | tr | tr | tr | tr | tr | tr | tr |
| 1147 | UNK 1147 | tr | tr | tr | tr | – | tr | tr | tr | tr | tr | tr | – | tr |
| 1152 | UNK 1152 | tr | tr | – | – | tr | – | tr | tr | tr | – | – | – | tr |
| 1156 | UNK 1156 | tr | tr | tr | tr | tr | tr | tr | tr | tr | tr | tr | – | – |
| 1162 | UNK 1162 | tr | – | – | – | 0.1 | – | tr | tr | tr | – | – | – | – |
| 1164 | UNK 1164_1 | tr | tr | tr | – | 0.1 | tr | 0.1 | 0.1 | 0.1 | tr | tr | tr | 0.1 |
| 1164 | UNK 1164_2 | tr | tr | tr | 0.2 | tr | tr | tr | tr | tr | tr | tr | tr | – |
| 1171 | UNK 1171 | tr | tr | tr | 0.1 | – | tr | tr | tr | – | tr | – | – | – |
| 1173 | UNK 1173_1 | tr | tr | – | – | – | tr | – | – | – | – | – | – | – |
| 1173 | UNK 1173_2 | tr | – | – | – | tr | – | tr | tr | tr | tr | – | – | – |
| 1181 | UNK 1181 | tr | tr | – | tr | tr | – | tr | tr | tr | – | – | – | – |
| 1188 | UNK 1188 | tr | tr | – | tr | – | tr | – | tr | – | – | tr | – | – |
| 1192 | UNK 1192 | tr | tr | – | – | tr | – | – | – | – | – | – | – | – |
| 1207 | UNK 1207_1 | tr | – | – | – | tr | tr | – | tr | tr | tr | tr | – | tr |
| 1207 | UNK 1207_2 | tr | tr | – | – | tr | tr | tr | tr | tr | tr | – | tr | – |
| 1222 | UNK 1222_1 | tr | – | – | – | tr | – | – | tr | tr | – | tr | tr | tr |
| 1222 | UNK 1222_2 | tr | tr | – | – | tr | – | – | tr | – | – | – | – | tr |
| 1225 | UNK 1225 | tr | tr | tr | tr | tr | tr | tr | tr | tr | tr | – | tr | – |
| 1240 | UNK 1240 | tr | tr | tr | tr | tr | tr | tr | tr | tr | tr | tr | tr | – |
| 1243 | UNK 1243 | tr | tr | – | tr | tr | tr | tr | tr | tr | tr | – | – | – |
| 1249 | UNK 1249 | tr | tr | – | – | tr | – | tr | tr | tr | tr | – | tr | tr |
| 1251 | UNK 1251 | tr | – | – | – | tr | – | tr | tr | tr | tr | tr | – | tr |
| 1260 | UNK 1260 | tr | tr | – | tr | – | tr | – | – | – | – | – | – | – |
| 1262 | UNK 1262 | tr | tr | – | – | tr | – | tr | tr | tr | – | tr | tr | tr |
| 1270 | UNK 1270 | – | – | – | – | – | – | – | tr | tr | – | – | tr | – |
| 1273 | UNK 1273 | tr | tr | tr | tr | – | tr | tr | tr | tr | – | – | – | – |
| 1274 | UNK 1274 | tr | tr | tr | tr | tr | tr | tr | tr | tr | tr | tr | tr | tr |
| 1279 | UNK 1279 | tr | tr | tr | 0.1 | 0.1 | tr | tr | tr | 0.1 | tr | tr | tr | tr |
| 1282 | UNK 1282 | tr | tr | – | tr | – | tr | – | tr | tr | – | – | – | tr |
| 1283 | UNK 1283_1 | tr | tr | tr | – | – | tr | tr | tr | tr | tr | tr | – | – |
| 1283 | UNK 1283_2 | tr | tr | – | tr | tr | – | tr | tr | tr | – | tr | – | – |
| 1289 | UNK 1289 | tr | tr | – | tr | tr | tr | tr | tr | tr | – | tr | tr | tr |
| 1292 | UNK 1292 | tr | tr | – | tr | tr | – | – | tr | tr | – | tr | – | – |

| | | | | | | | | | | | | | | |
|---|---|---|---|---|---|---|---|---|---|---|---|---|---|---|
| 1300 | UNK 1300 | tr | tr | – | – | – | – | – | tr | tr | tr | – | – | – |
| 1337 | UNK 1337 | tr | tr | tr | tr | tr | tr | tr | tr | tr | tr | tr | tr | – |
| 1339 | UNK 1339 | tr | tr | tr | tr | tr | tr | tr | tr | tr | tr | tr | tr | tr |
| 1341 | UNK 1341 | tr | tr | – | – | tr | – | tr | tr | tr | tr | tr | tr | tr |
| 1347 | UNK 1347 | tr | 0.1 | tr | 0.1 | 0.2 | tr | 0.1 | tr | tr | tr | tr | tr | 0.1 |
| 1349 | UNK 1349 | tr | tr | tr | 0.1 | tr | tr | 0.1 | tr | tr | 0.2 | tr | tr | – |
| 1355 | UNK 1355 | tr | tr | tr | – | – | tr | tr | tr | tr | tr | tr | tr | – |
| 1359 | UNK 1359 | tr | tr | – | – | – | – | – | tr | tr | tr | tr | – | – |
| 1367 | UNK 1367_1 | 0.1 | 0.2 | tr | 0.1 | 0.2 | tr | 0.1 | 0.2 | 0.1 | 0.1 | 0.4 | 0.3 | 0.4 |
| 1367 | UNK 1367_2 | tr | tr | tr | tr | tr | tr | tr | tr | tr | tr | tr | tr | tr |
| 1371 | UNK 1371 | tr | tr | tr | tr | tr | tr | tr | tr | tr | tr | tr | tr | tr |
| 1378 | UNK 1378 | tr | 0.1 | tr | tr | tr | tr | tr | tr | tr | 0.1 | 0.1 | 0.1 | tr |
| 1381 | UNK 1381 | tr | tr | tr | tr | 0.1 | tr | tr | 0.1 | tr | tr | 0.2 | 0.1 | 0.2 |
| 1388 | UNK 1388 | tr | tr | tr | tr | tr | tr | tr | tr | tr | tr | tr | tr | tr |
| 1391 | UNK 1391 | 0.1 | 0.1 | 0.1 | 0.1 | tr | 0.1 | 0.1 | 0.1 | 0.1 | 0.1 | 0.1 | 0.1 | 0.1 |
| 1392 | UNK 1392 | tr | tr | – | tr | – | – | tr | tr | tr | tr | – | – | – |
| 1394 | **UNK 1394** | 0.2 | 0.2 | tr | 0.1 | 0.6 | 0.2 | 0.2 | 0.1 | 0.1 | 0.1 | 1.1 | 0.2 | 0.1 |
| 1403 | UNK 1403 | tr | 0.1 | tr | tr | tr | tr | tr | tr | tr | 0.1 | 0.1 | 0.1 | tr |
| 1409 | **UNK 1409_1** | 0.2 | 0.2 | tr | 0.4 | 0.4 | 0.1 | 0.1 | 0.2 | 0.2 | 0.1 | 0.6 | 0.2 | 1.1 |
| 1409 | UNK 1409_2 | tr | tr | – | – | – | – | – | – | – | – | – | – | – |
| 1415 | **UNK 1415** | 3.7 | 3.9 | 1.7 | 2.3 | 7.3 | 3.7 | 3.4 | 3.8 | 3.1 | 2.8 | 10.4 | 2.3 | 11.3 |
| 1416 | UNK 1416 | tr | tr | tr | tr | – | – | – | tr | tr | tr | tr | tr | – |
| 1421 | UNK 1421 | tr | tr | tr | – | tr | – | tr | tr | tr | – | tr | – | – |
| 1424 | UNK 1424 | tr | tr | tr | tr | tr | tr | tr | tr | tr | tr | tr | tr | tr |
| 1425 | UNK 1425 | 0.1 | 0.2 | 0.1 | 0.1 | 0.1 | 0.1 | 0.1 | 0.2 | 0.1 | 0.2 | 0.2 | 0.3 | 0.2 |
| 1426 | UNK 1426 | tr | tr | tr | tr | tr | tr | tr | tr | tr | – | tr | tr | tr |
| 1427 | UNK 1427 | tr | tr | – | – | – | tr | – | tr | tr | tr | – | tr | – |
| 1431 | UNK 1431 | tr | tr | tr | tr | tr | tr | tr | tr | tr | tr | tr | tr | tr |
| 1438 | UNK 1438_1 | 0.1 | 0.1 | tr | tr | tr | tr | 0.1 | tr | tr | 0.1 | tr | 0.1 | tr |
| 1438 | UNK 1438_2 | tr | tr | tr | tr | tr | tr | tr | tr | tr | – | – | tr | – |
| 1440 | UNK 1440 | 0.1 | 0.1 | 0.1 | 0.1 | 0.2 | tr | 0.1 | 0.1 | 0.1 | 0.2 | 0.2 | 0.1 | 0.1 |
| 1441 | UNK 1441 | tr | tr | tr | tr | tr | tr | tr | tr | tr | 0.1 | 0.1 | 0.1 | tr |
| 1447 | UNK 1447 | tr | tr | tr | tr | tr | tr | – | tr | tr | tr | – | tr | – |
| 1453 | UNK 1453 | tr | tr | tr | tr | tr | tr | tr | tr | tr | tr | tr | tr | tr |
| 1457 | UNK 1457 | 0.2 | 0.3 | 0.2 | 0.1 | tr | 0.1 | 0.2 | 0.2 | 0.1 | 0.3 | 0.2 | 0.3 | tr |
| 1459 | UNK 1459 | tr | tr | tr | tr | tr | tr | 0.1 | tr | tr | tr | 0.1 | tr | 0.1 |
| 1460 | UNK 1460 | tr | tr | – | – | – | – | tr | tr | tr | tr | tr | tr | – |
| 1465 | UNK 1465 | tr | tr | – | tr | 0.1 | tr | tr | tr | tr | tr | tr | tr | tr |
| 1466 | UNK 1466_1 | 0.2 | 0.3 | 0.1 | 0.1 | 0.1 | 0.2 | 0.2 | 0.2 | 0.1 | 0.2 | 0.3 | 0.4 | 0.2 |
| 1466 | UNK 1466_2 | tr | tr | tr | tr | – | tr | tr | tr | tr | tr | tr | tr | tr |
| 1469 | UNK 1469_1 | tr | tr | tr | – | tr | tr | tr | tr | tr | tr | tr | tr | tr |
| 1469 | UNK 1469_2 | tr | tr | – | tr | tr | 0.1 | tr | – | – | – | – | – | – |
| 1472 | UNK 1472 | 0.1 | 0.2 | tr | 0.1 | tr | tr | 0.1 | 0.1 | tr | 0.1 | 0.1 | 0.1 | 0.1 |
| 1477 | UNK 1477 | tr | tr | tr | – | – | – | tr | tr | tr | tr | tr | tr | – |
| 1481 | UNK 1481 | 0.4 | 0.5 | 0.3 | 0.1 | 0.3 | 0.4 | 0.6 | 0.2 | 0.1 | 0.2 | 0.4 | tr | 0.3 |
| 1483 | UNK 1483 | tr | tr | – | tr | tr | – | tr | tr | tr | tr | tr | tr | tr |
| 1484 | UNK 1484 | tr | tr | tr | – | – | – | – | tr | tr | tr | – | tr | – |
| 1486 | UNK 1486 | tr | tr | tr | – | tr | tr | tr | tr | tr | tr | tr | – | tr |

| | | | | | | | | | | | | | | |
|---|---|---|---|---|---|---|---|---|---|---|---|---|---|---|
| 1488 | UNK 1488 | tr | tr | tr | tr | tr | 0.1 | 0.1 | tr | tr | tr | tr | tr | tr |
| 1492 | **UNK 1492** | 1.7 | 2.7 | 1.4 | 0.3 | 0.5 | 0.5 | 1.8 | 1.2 | 1 | 1.4 | 1.6 | 3.1 | 0.6 |
| 1496 | UNK 1496 | tr | 0.1 | tr | tr | tr | tr | 0.1 | tr | tr | tr | 0.1 | tr | tr |
| 1503 | **UNK 1503** | 0.8 | 0.9 | 0.2 | 0.4 | 0.8 | 0.6 | 1 | 0.2 | 0.2 | 0.1 | 0.4 | 0.2 | 0.3 |
| 1507 | UNK 1507_1 | tr | tr | – | tr | tr | tr | tr | tr | tr | tr | tr | tr | tr |
| 1507 | UNK 1507_2 | tr | tr | – | tr | – | – | – | – | – | – | – | – | – |
| 1510 | UNK 1510 | tr | tr | – | – | – | tr | – | tr | tr | tr | tr | tr | – |
| 1512 | UNK 1512 | tr | tr | tr | – | – | – | tr | tr | tr | tr | tr | – | – |
| 1517 | UNK 1517 | tr | tr | tr | tr | tr | – | tr | tr | tr | tr | tr | tr | tr |
| 1518 | UNK 1518 | 0.1 | 0.2 | 0.1 | 0.1 | tr | 0.1 | 0.1 | 0.1 | 0.1 | 0.2 | 0.1 | 0.1 | 0.1 |
| 1521 | UNK 1521_1 | tr | – | – | – | tr | tr | – | tr | tr | tr | tr | – | – |
| 1521 | UNK 1521_2 | tr | tr | tr | tr | – | 0.4 | tr | tr | tr | tr | tr | 0.1 | tr |
| 1523 | UNK 1523 | tr | tr | tr | – | – | – | – | tr | tr | – | – | – | – |
| 1524 | **UNK 1524** | 0.7 | 1 | 1.3 | 0.2 | 0.2 | 0.8 | 0.7 | 0.4 | 0.2 | 0.4 | 0.5 | 0.9 | 0.5 |
| 1526 | UNK 1526_1 | tr | tr | – | tr | tr | – | tr | tr | tr | tr | tr | tr | tr |
| 1526 | UNK 1526_2 | tr | tr | – | – | tr | – | – | tr | tr | tr | – | – | – |
| 1527 | UNK 1527 | tr | tr | – | tr | – | tr | – | tr | tr | tr | tr | – | tr |
| 1532 | UNK 1532 | tr | tr | tr | – | tr | tr | tr | tr | tr | tr | tr | tr | tr |
| 1541 | UNK 1541 | 0.1 | 0.1 | 0.1 | 0.1 | 0.1 | 0.1 | 0.1 | 0.1 | tr | 0.1 | 0.1 | 0.1 | 0.1 |
| 1553 | UNK 1553 | tr | tr | tr | 0.1 | tr | 0.1 | 0.1 | tr | tr | tr | tr | tr | tr |
| 1556 | UNK 1556 | tr | tr | – | – | tr | tr | tr | tr | tr | – | tr | tr | – |
| 1559 | UNK 1559_1 | tr | tr | – | – | – | – | tr | tr | tr | – | tr | – | tr |
| 1559 | UNK 1559_2 | tr | tr | – | – | tr | – | – | tr | tr | – | tr | – | – |
| 1560 | UNK 1560 | – | – | – | – | – | – | – | tr | tr | tr | – | tr | – |
| 1563 | UNK 1563 | tr | tr | tr | tr | tr | tr | tr | tr | tr | – | – | tr | tr |
| 1576 | UNK 1576 | tr | tr | tr | – | tr | tr | – | tr | tr | tr | tr | – | tr |
| 1582 | UNK 1582 | tr | 0.1 | tr | tr | tr | tr | tr | tr | tr | tr | tr | tr | tr |
| 1584 | UNK 1584 | tr | tr | tr | tr | 0.1 | tr | 0.1 | tr | tr | tr | tr | tr | 0.1 |
| 1590 | UNK 1590 | tr | tr | – | tr | tr | tr | tr | tr | tr | tr | tr | tr | tr |
| 1595 | UNK 1595 | tr | tr | tr | tr | tr | 0.1 | tr | tr | tr | tr | tr | tr | tr |
| 1596 | UNK 1596 | tr | tr | tr | tr | tr | tr | tr | tr | tr | tr | – | – | – |
| 1599 | UNK 1599 | tr | tr | tr | tr | tr | – | tr | tr | tr | tr | 0.1 | tr | tr |
| 1601 | UNK 1601 | tr | tr | – | tr | tr | tr | tr | tr | tr | tr | tr | tr | tr |
| 1603 | UNK 1603 | tr | tr | – | – | tr | – | tr | tr | – | – | – | – | tr |
| 1607 | UNK 1607 | tr | tr | tr | tr | 0.2 | tr | tr | tr | tr | tr | 0.1 | tr | tr |
| 1609 | UNK 1609 | tr | tr | tr | tr | – | – | tr | tr | tr | tr | tr | – | – |
| 1610 | UNK 1610 | tr | – | tr | – | – | tr | – | – | – | – | – | – | – |
| 1614 | UNK 1614 | tr | tr | tr | – | – | – | – | tr | tr | tr | tr | tr | – |
| 1616 | UNK 1616 | 0.1 | 0.1 | tr | tr | 0.1 | 0.1 | 0.2 | tr | tr | 0.1 | 0.1 | tr | 0.1 |
| 1617 | UNK 1617 | tr | tr | tr | – | tr | tr | tr | tr | tr | tr | tr | tr | tr |
| 1625 | UNK 1625 | tr | tr | – | – | tr | – | tr | tr | tr | – | tr | – | – |
| 1629 | UNK 1629 | tr | tr | tr | – | – | – | tr | tr | tr | tr | – | tr | – |
| 1633 | UNK 1633 | tr | tr | – | – | tr | – | – | tr | tr | tr | – | – | – |
| 1640 | UNK 1640 | tr | tr | tr | tr | tr | 0.1 | tr | tr | tr | tr | tr | tr | – |
| 1643 | UNK 1643 | tr | tr | tr | tr | tr | tr | tr | tr | tr | tr | tr | tr | tr |
| 1650 | UNK 1650 | tr | tr | tr | tr | tr | 0.1 | 0.1 | tr | tr | tr | tr | 0.1 | tr |
| 1654 | UNK 1654 | tr | tr | tr | tr | tr | tr | tr | tr | tr | tr | tr | tr | tr |
| 1658 | UNK 1658 | 0.1 | 0.1 | tr | tr | tr | 0.1 | 0.1 | tr | tr | 0.1 | 0.1 | tr | 0.2 |

| | | | | | | | | | | | | | | |
|---|---|---|---|---|---|---|---|---|---|---|---|---|---|---|
| 1660 | UNK 1660 | tr | tr | – | 0.1 | tr | tr | – | tr | tr | tr | tr | tr | tr |
| 1667 | UNK 1667 | tr | tr | tr | tr | tr | tr | 0.1 | tr | tr | tr | tr | – | tr |
| 1670 | UNK 1670 | tr | tr | – | tr | – | – | – | tr | tr | tr | – | tr | – |
| 1671 | UNK 1671 | tr | tr | tr | – | tr | tr | tr | tr | – | – | tr | – | – |
| 1672 | UNK 1672 | tr | tr | – | – | – | – | – | tr | tr | – | – | – | – |
| 1675 | UNK 1675 | tr | tr | – | tr | – | tr | – | tr | tr | – | – | – | – |
| 1677 | UNK 1677 | tr | tr | tr | – | – | tr | tr | tr | tr | tr | tr | – | – |
| 1699 | **UNK 1699**[†] | 3.6 | 4.1 | 3 | 0.5 | 1.8 | 3.2 | 5.2 | 1.3 | 1.1 | 1.6 | 2 | 0.8 | 3.2 |
| 1707 | UNK 1707 | tr | tr | – | tr | – | – | – | – | – | – | – | – | – |
| 1710 | UNK 1710 | tr | tr | – | – | tr | tr | tr | tr | tr | tr | tr | – | – |
| 1711 | UNK 1711 | tr | tr | – | tr | tr | tr | tr | tr | tr | tr | tr | tr | tr |
| 1716 | UNK 1716 | tr | tr | tr | tr | tr | tr | tr | tr | tr | tr | – | tr | – |
| 1735 | UNK 1735 | tr | tr | – | – | tr | – | tr | tr | tr | tr | – | – | – |
| 1749 | UNK 1749 | tr | tr | – | – | tr | – | – | tr | – | – | tr | – | – |
| 1757 | UNK 1757_1 | 0.2 | 0.2 | 0.1 | 0.2 | 0.2 | 0.6 | 0.5 | 0.1 | 0.1 | 0.4 | 0.1 | tr | tr |
| 1757 | UNK 1757_2 | tr | tr | tr | – | tr | – | tr | tr | tr | tr | tr | tr | tr |
| 1773 | UNK 1773 | tr | tr | – | – | tr | – | tr | tr | tr | tr | – | tr | – |
| 1794 | UNK 1794 | 0.1 | 0.1 | 0.3 | 0.1 | 0.1 | 0.2 | 0.3 | tr | 0.1 | tr | tr | 0.1 | tr |
| 1819 | UNK 1819 | tr | tr | – | – | tr | – | tr | – | – | – | – | – | – |
| 1854 | UNK 1854 | tr | tr | tr | 0.1 | 0.1 | tr | 0.2 | tr | tr | tr | 0.3 | tr | tr |
| 1875 | UNK 1875 | tr | tr | – | – | – | tr | – | tr | – | – | tr | – | tr |

[§]Synthetic (+)-$\alpha$- and (+)-$\beta$-Citronellene coeluted with natural detected $\alpha$- and $\beta$-Citronellene on a chiral column (MEGA-DEX DMT Beta SE, 30m × 0.25mm ID, 0.23µm film) (Gfrerer *et al*. unpublished)

[†]Also detected by (*56*) where it was referred to as “unknown 53”
